# Mosaic changes to the global transcriptome in response to inhibiting ribosome formation versus inhibition of ribosome function

**DOI:** 10.1101/2020.10.15.341230

**Authors:** Md Shamsuzzaman, Nusrat Rahman, Brian Gregory, Vincent M Bruno, Lasse Lindahl

## Abstract

Cell fate is susceptible to several internal and external stresses. Stress resulting from mutations in genes for ribosomal proteins and assembly factors leads to many congenital diseases, collectively called ribosomopathies. Even though such mutations all depress the cell’s protein synthesis capacity, they are manifested in many different phenotypes. This prompted us to use *Saccharomyces cerevisiae* to explore whether reducing the protein synthesis capacity by different mechanisms result in the same or different changes to the global transcriptome. We have compared the transcriptome after abolishing the assembly of new ribosomes and inhibiting the translocation of ribosomes on the mRNA. Our results show that these alternate obstructions generate different mosaics of expression for several classes of genes, including genes for ribosomal proteins, mitotic cell cycle, cell wall synthesis, and protein transport.

## Introduction

The formation of eukaryotic ribosomes is a complex process, largely conserved from *Saccharomyces cerevisiae* (yeast) to humans, involving the synthesis of two rRNA transcripts, 79-80 ribosomal proteins (r-proteins), and over 250 assembly factors (Bassler and Hurt 2019; Bohnsack and Bohnsack 2019; Klinge and Woolford 2019). This process is very resource-intensive and mechanisms have evolved to adjust the ribosome production that minimizes the number of idle ribosomes and assures ribosome homeostasis (Kief and Warner 1981). This regulation includes a variety of responses to environmental stress such as nutrient starvation, exposure to oxidative chemicals, and changes in temperature or osmotic value that down-regulate r-protein genes together with a large number of other genes while up-regulating genes encoding proteins needed to adapt to the stress condition (Gasch et al. 2000; Ho and Gasch 2015; Piazzi et al. 2019).

Yeast is the best-investigated model for the regulation of eukaryotic ribosome synthesis. The transcription of r-protein genes by RNA polymerase II is regulated by TORC1, which under favorable conditions stimulates the binding of the Fhl1, Ifh1, and Rap1 or Abf1 transcription factors upstream of the genes (Martin et al. 2004; Fermi et al. 2017). During stress, TOR is inactivated resulting in replacing Ifh1 with the repressor Rfc1. Recent evidence shows that these signaling pathways originate, at least in part, from interference with normal ribosome function, which obliterates ribosome homeostasis. One such regulatory process originates from ribosomal collisions on the mRNA that activate the kinase Gcn2 to phosphorylate eIF2α triggering inhibition of translation initiation (Wu et al. 2019; Juszkiewicz et al. 2020; Wu et al. 2020) and favors the expression of a subset of genes through ribosome interaction with upstream open reading frames (Hinnebusch et al. 2016; Pakos-Zebrucka et al. 2016). During extreme stress, another kinase, ZAKα, phosphorylates map kinases resulting in apoptosis and cell death (Vind et al. 2020; Wu et al. 2020). Another pathway limiting expression of r-protein genes is actuated by obstruction of ribosome assembly, which causes accumulation of insoluble aggregates of ribosomal and other proteins including Ifh1 (Albert et al. 2019a).

Mutations in numerous genes for r-proteins or assembly factors produce stress leading to a variety of congenital human diseases, collectively called “ribosomopathies” (Armistead and Triggs-Raine 2014; Aubert et al. 2018). Furthermore, several r-proteins have been identified as cancer drivers (Fancello et al. 2017). The suggested roots for these calamities include decreased ribosome production with an ensuing decrease in translation capacity, the formation of “specialized ribosomes” with altered preferences for specific mRNAs, accumulation of free r-proteins that interact with p53 and other cell cycle regulators, and changes to specific metabolic pathways (Deisenroth and Zhang 2011; Kondrashov et al. 2011; Bursac et al. 2014; James et al. 2014; Raiser et al. 2014; Zhou et al. 2015; Mills and Green 2017; Farley-Barnes et al. 2019; Maitra et al. 2020). One of the standing puzzles of ribosomopathies is that although mutations in all r-protein genes decrease the rate of ribosome formation, mutations in different ribosomal genes generate a variety of distinct disease phenotypes, and, vice versa, mutations in different r-protein genes can cause the same disease, e.g. Diamond Black anemia (Ulirsch et al. 2018).

Typically, investigations of cell stress focus on the short-term changes in signaling pathways and gene expression immediately following the onset of the stress. However, the ultimate cell fate depends as much on secondary changes in gene expression evolving during sustained stress. To gain insight into the long-term effects of permanently assaulting the cell’s capacity for protein synthesis, we asked if mutations limiting protein synthesis in different ways generate the same long-term stress response. To this end, we used conditional *Saccharomyces* cerevisiae (yeast) mutants to compare the evolving changes to the transcriptome after ceasing the synthesis of r-protein uL4 (“ribosomal stress”) or abolishing the synthesis of Translation Elongating Factor eEF3 (“translation stress”). Both these procedures lower the translation capacity, but in completely different ways (Fig 1A). Ribosomal stress abrogates ribosomal assembly and lowers the number of ribosomes without inhibiting ribosome function, while translation stress prevents ribosomal translocation on the mRNA without changing the total number of ribosomes or the number of ribosomes on polysomes (Shamsuzzaman et al. 2017; Gregory et al. 2019).

**Figure 1.**
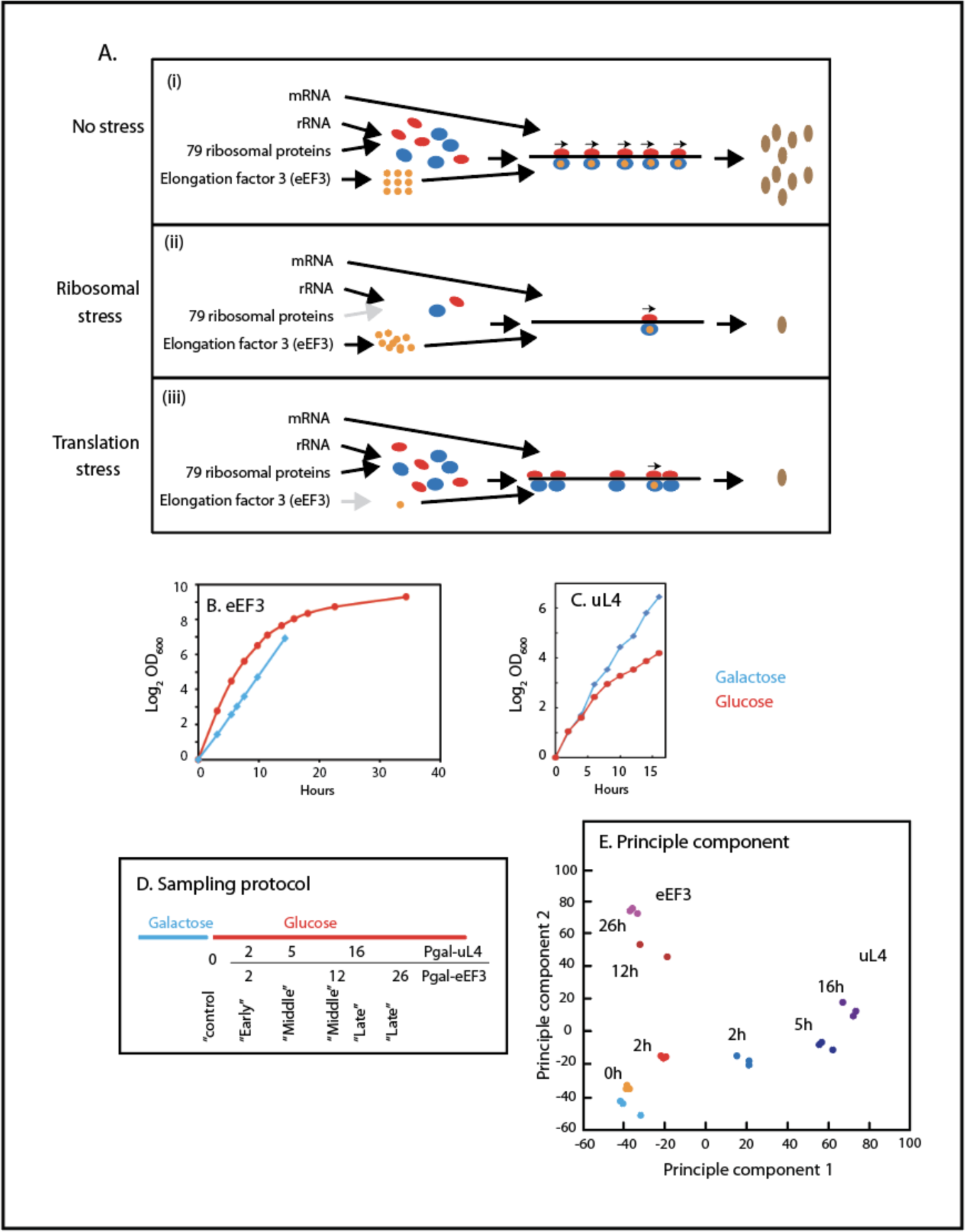
Experimental design, growth curves, and Principal Component Analysis. (**A**) Experimental design. Top. Uninhibited protein synthesis. 40S and 60S ribosomal subunits in red and blue color, respectively, eEF3 in orange, and completed proteins in brown. Arrows above the 80S ribosomes on the mRNA indicate translocation competent ribosomes. Middle. Ribosomal stress: ribosome assembly is inhibited by repression of uL4 synthesis, resulting in reduced numbers of ribosomes, but the remaining ribosomes are translocation competent because the number of eEF3 molecules is not decreased. Bottom. Translation stress: the number of ribosomes in unchanged, but eEF3 is depleted. Ribosomes become translocation incompetent as the number of eEF3 molecules decreases. A mixture of translocation competent and translocation incompetent ribosomes on an mRNA may lead to ribosome collisions. (**B** and **C**). Growth curves of Pgal-eEF3 and Pgal-uL4, respectively, grown in galactose (blue curves) and after a shift from galactose to glucose (orange curves). (**D**) The schema for the sampling of cultures. The sampling times were designed to match the growth rates of the two strains at the time of withdrawing the control, early, middle and late samples. The timing of each sample is indicated. (**E**) Principal component analysis of the RNA-seq data in the two strains in samples taken at the indicated times. Results from Pgal-uL4 are shown by colors in the blue spectrum and Pgal-eEF3 results are shown by colors in the yellow-to-red spectrum.

Our results show that ribosomal and translation stresses generate very different transcriptomes. Genes under several GO terms respond differently to each of the two types of stress, while genes under other GO terms respond similarly. It is especially remarkable that r-protein genes are regulated in opposite directions, up during ribosomal stress, down during translation stress. Thus, there is not a common stress response to lowering the translation capacity, but ribosomal and translation stress generate unique mosaic transcriptomes.

## Results

### Establishing ribosome and translation stress

To establish the ribosomal and translation stress conditions, we used *S. cerevisiae* strains in which the only gene for r-protein uL4 (Pgal-uL4) or Translation Elongation Factor eEF3 (Pgal-eEF3) is expressed from the *GAL1/10* galactose promoter (Table S1). Shifting these strains from galactose to glucose medium essentially ceases the synthesis of the protein under gal control. Abolishing the synthesis of uL4, or any other r-protein blocks ribosome assembly and reduces the ribosome number without affecting ribosome function (Gregory et al. 2019) while repressing eEF3 synthesis inhibits the ribosome without changing the ribosome content (Fig 1A) (Shamsuzzaman et al. 2017).

The growth curve for Pgal-eEF3 shows a typical “shift-up response”, i.e. an increase of the growth rate immediately after the shift to glucose medium due to an upsurge in the differential rate of ribosome formation (Fig 1B) (Kjeldgaard et al. 1958; Kief and Warner 1981). However, no shift-up was seen in the Pgal-uL4 growth curve, as expected, since ribosome production is blocked, preventing an increase in ribosome number (Fig 1C). The effect of the shift-up on gene expression stabilizes within an hour after the shift (Kief and Warner 1981), leading us to reason that the effect of the two stress forms on the transcriptome can be compared after 2 hours.

### General trends of the transcriptome

We collected samples from triplicate cultures of Pgal-uL4 and Pgal-eEF3 at time points chosen to roughly match the growth rates of the two strains at the time of sampling. We refer to the samples as “control”, “early”, “mid”, and “late” stress samples (Fig 1D). One of the middle Pgal-eEF3 samples was discarded due to poor RNA quality. RNA-seq data were mapped to the genome of the parent strain (BY4741) (Table S2), yielding 2.3-3.1*10^7^ and 2.5-3.3*10^7^ reads from each sample of Pgal-uL4 and Pgal-eEF3, respectively.

The replicate samples cluster tightly in principal component analysis (PCA), indicating high reproducibility (Fig 1E). Moreover, the results from the control samples from the two strains cluster together suggesting a strong similarity between the transcriptomes in the two strains during uninhibited growth. After the induction of stress, the results move along different paths in the PCA space, indicating that the transcriptome evolves differently during the two stress forms. Except for the late Pgal-uL4 samples, the expression of over 40% of the genes changed less than 2.8-fold up or down (−1.5< log_2_ of expression<1.5) (Fig S1A). Only 29% of the genes in Pgal-uL4 and 16% of the genes in Pgal-eEF3 changed more than 2-fold at all times, a trend manifested for both up% and down-regulated genes (Fig S1B-D). Moreover, the differential trend of the transcriptome seen in the PCA (Fig 1E) was confirmed by fact that only 496 of 1884 (26%) DEG, defined as ≥2-fold changes up or down, are shared between late samples from Pgal-uL4 and Pgal-eEF3 (Fig S1E) with similar trends for both up- and down-regulated genes. Overall, the general analysis indicates that the transcriptome responds differently during ribosomal and translation stress.

We identified the genes with the most different trends during the two stress forms by ranking of gene pairs (Heinaniemi et al. 2013). The genes most likely to be expressed differently in the two strains, on average, over the three time points and are referred to as “the top differentiating genes”. Heat maps for genes on the top differentiating genes are shown in Fig 2. The list includes several genes for the oxidative stress response, cell wall synthesis, and amino acid biosynthesis. Comparing the expression data by enrichment analysis and Wilcoxon rank test revealed GO-terms mitotic cell cycle, protein transport, and fungal type cell wall biogenesis on the list of the top differentiating genes. Finally, the function of the genes with the most different expression was identified by overrepresentation analysis of these genes using CPDB yeast (http://cpdb.molgen.mpg.de/YCPDB). GO terms significantly overrepresented at the top of this list include mostly ribosome biogenesis, vesicles, fungal type cell wall biogenesis. Furthermore, the stress responses would be most obvious if the expression trend changed sign after the initial response caused by the shift of growth medium seen in the early samples. Accordingly, queried the list of most different rank-sum for genes that initially show increased expression, but changed to a negative trend after the early sample was taken, or switched in the opposite direction from negative to a positive trend. Most of the top 50 genes on this list were r-protein genes (not shown).

**Figure 2.**
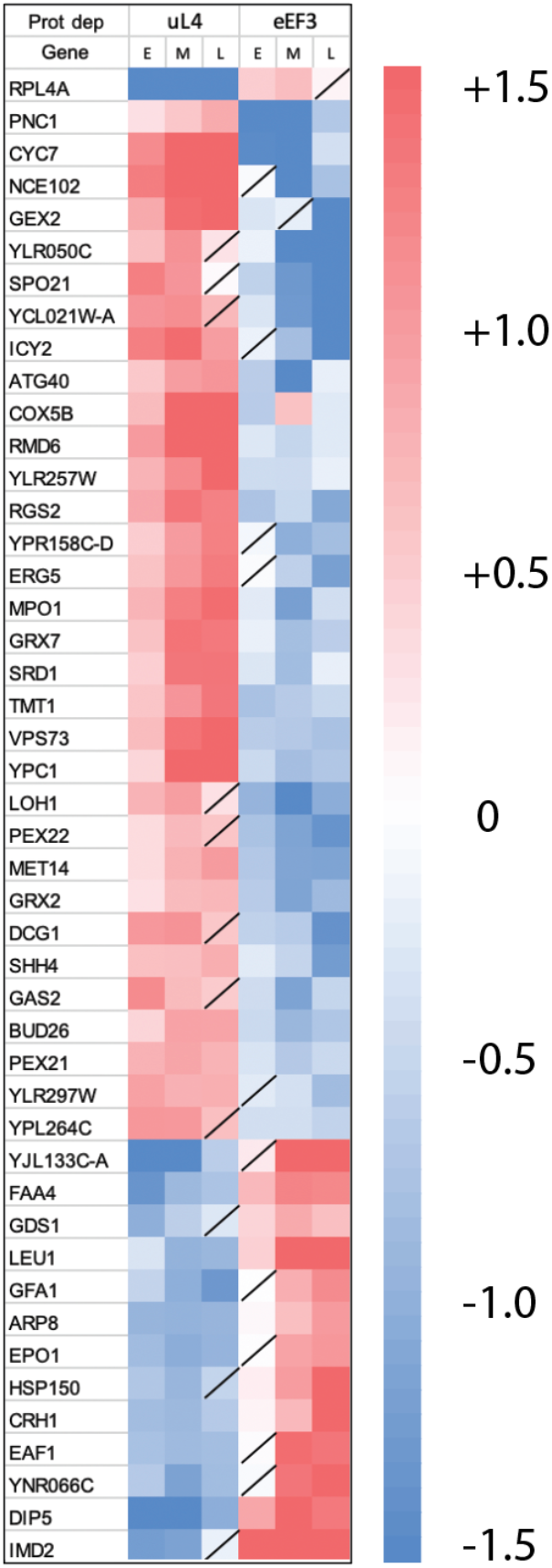
Genes expressed most differently after repressing uL4 or eEF3 synthesis. Heatmap of the expression of top 46 reading frames with the highest absolute cumulative rank differences in expression between ribosomal and translation stress established by the repression of uL4 and eEF3 synthesis. E, M, and L indicate early, middle, and late samples (Fig 1). See Materials and Methods for the algorithm used to calculate the rank differences. Reading frames with an FDR >0.05 are indicated with a line across the field on the heating map. All others have an FDR≤0.05

### Dissecting the expression of genes in the categories identified in the general trend analysis

#### Genes for ribosome biogenesis

The heat diagram and mean gene expression change for r-protein genes (Fig 3A and 3B (i)) show that repression of uL4 synthesis produces an initial downward trend followed by a switch to a monotonous increase after 2 hours. Repression of eEF3 synthesis results in the opposite pattern: an initial upsurge as expected from the nutritional shift-up followed by a decrease throughout the rest of the experiment. Fig 3A-B(i) shows results for genes for which at least 5 of the 6 time points have an FDR ≤0.05, but as seen in Fig S2, even genes with FDR>0.05 follow the same pattern, with few exceptions.

**Figure 3.**
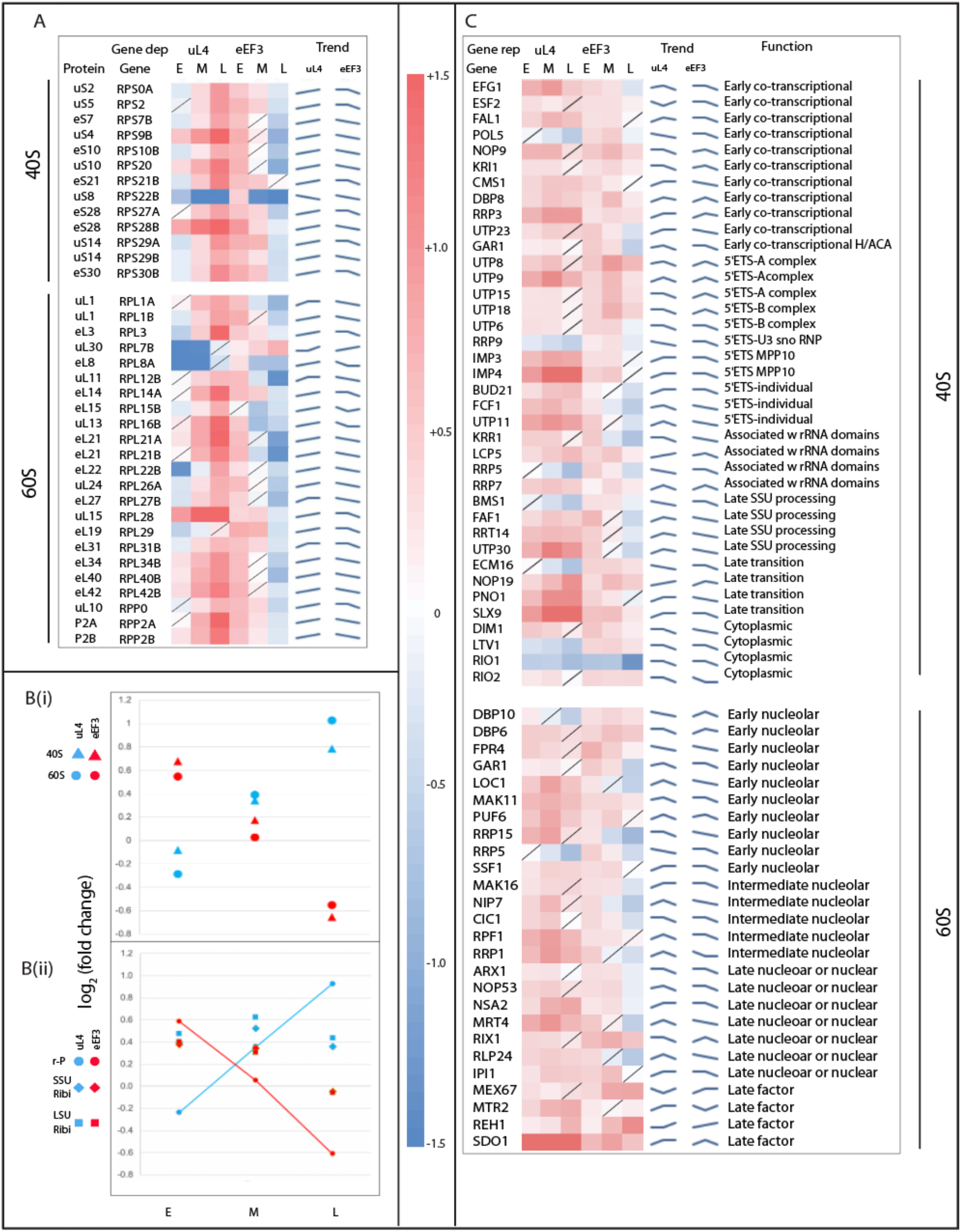
Expression of genes for cytoplasmic ribosomal proteins and ribosomal assembly factors. (**A**) Heatmaps for expression of the 60S and 40S ribosomal protein genes. The nomenclature for genes follows the standards of the Saccharomyces Genome Database (https://www.yeastgenome.org/). The names of proteins encoded by the r-protein genes are indicated following the current nomenclature of ribosomal proteins (Ban et al. 2014). (**B**) (i) Scatterplots for the mean log_2_ changes (LFC) in the expression of 60S and 40S ribosomal protein genes. (ii) Scatter plot for LFC of ribosomal assembly genes (Ribi) for each of the ribosomal subunits as well as the average expression of r-protein genes from both subunits. (**C**) Heat diagrams for expression of assembly factors for cytoplasmic ribosomes. “Trend lines” are calculated by the “Sparkline” algorithm in Excel.

Fig 3C shows the heat diagrams for ribosomal assembly protein genes (Ribi) listed in the order of the specific reactions supported by each factor (Klinge and Woolford 2019). Fig 3B(ii) shows the mean gene expression change for the 40S- and 60S-specific Ribi genes as well as the mean gene change for the average of r-protein genes for both subunits. The mean expression changes for assembly factors for both 40S and 60S Ribi genes differ from the patterns observed for r-protein genes. All Ribi genes initially increase 40-50%, but the 60S Ribi genes decrease slightly during late phases of both types of stress, while the 40S Ribi genes do not.

All genes for mitochondrial r-proteins and translation factors are repressed around 2-fold, on average, after shifting either Pgal-uL4 or Pgal-eEF3 to glucose medium (Fig S3A-C). This was expected since mitochondrial ribosomes only translate proteins for the cytochrome oxidative complex, which are also heavily repressed after the shift to glucose medium (Fig S3D) in agreement with previous reports (Costanzo and Fox 1990; Ulery et al. 1994; Ott et al. 2016).

#### Genes involved in cell cycle

We have previously shown that post-mitotic cell separation is inhibited by both translation and ribosomal stress as indicated by the progressive accumulation of daughter-mother cell complexes (Shamsuzzaman et al. 2017). This concords with a waning expression of daughter specific genes, including genes encoding three glucanase-like proteins involved in degrading the cell wall from the daughter side (Colman-Lerner et al. 2001; Weiss 2012) (Fig 4A). These and other daughter-specific genes trending downward (Fig S4A) are repressed by a Crz1 cascade. The actuation of such a cascade during both stress forms was confirmed by the upward trend of genes that are positively regulated by Crz1 after the repression of both uL4 and eEF3 synthesis (Fig. S4A). These results are consistent with the model that the cell cycle is arrested by Crz activity during both types of stress (see also transcription factor analysis below).

**Figure 4.**
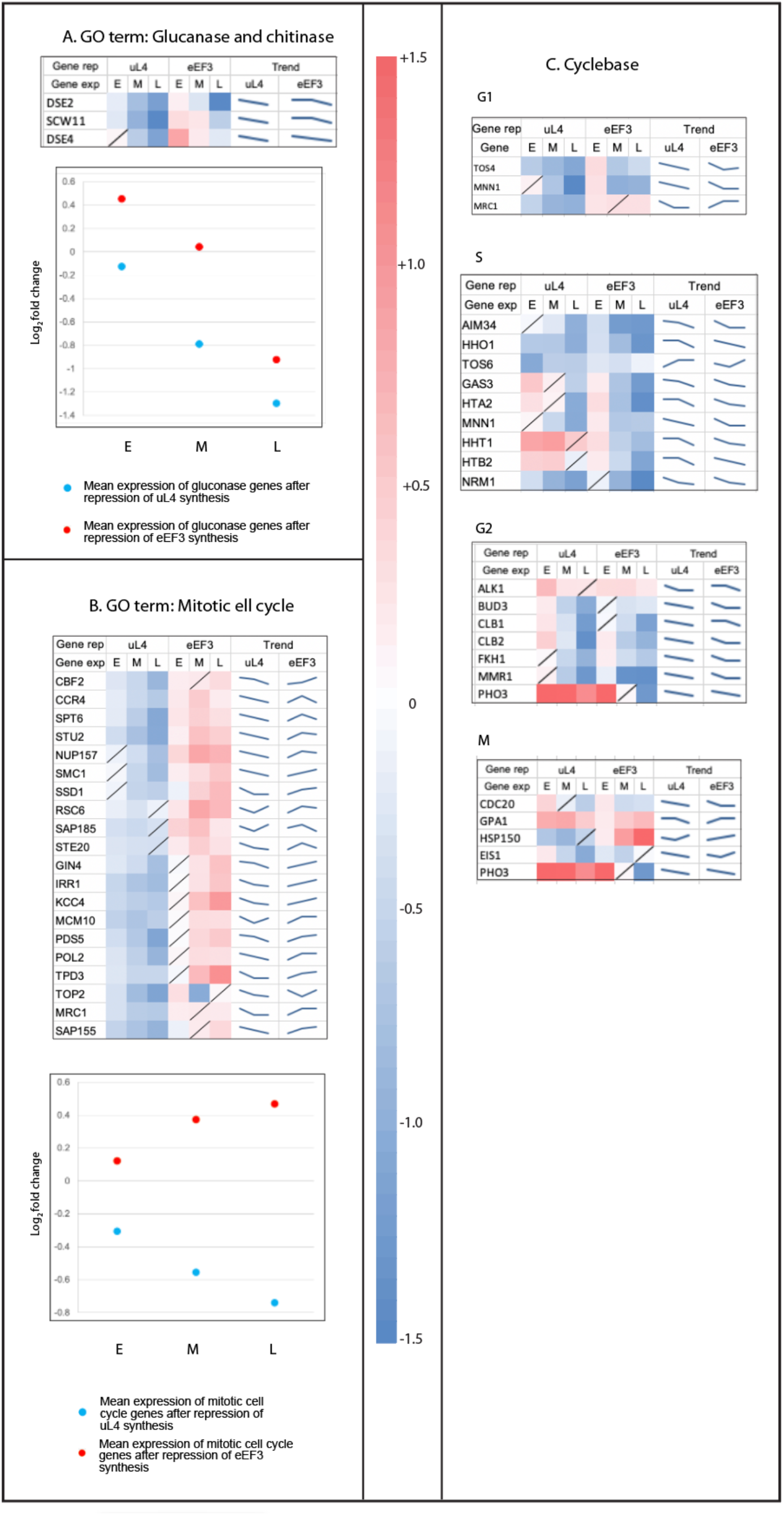
Expression of cell cycle genes. (A) Heatmap and scatterplot for the change in expression of genes cataloged under the GO-term “Mitotic Cell Cycle”. (**B**) Heatmaps for the changes in expression of genes with expression identified in the Cyclebase database ((http://www.cyclebase.org/CyclebaseSearch) as specific to the indicated phases of the cell cycle. (**C**) Heatmap for the expression of glucanases. For FDR statistics and Trend lines, see legends to Fig 2 and 3.

Interestingly, genes under the GO-term “Mitotic Cell Cycle” trend negatively during ribosomal stress, but positively during translational stress (Fig 4B). We also used Cyclebase (http://www.cyclebase.org/) to identify genes that are not included in the Mitotic Cell Cycle GO-term, but whose expression peaks specifically during different phases of the cell cycle (see) (Fig 4C). However, none of these genes were consistently repressed during both types of stress.

#### Other genes with different responses during translation and ribosomal stress

As seen from Fig 5A-B, genes under the GO-terms Chitin Biogenesis and Fungal Cell Wall Biogenesis trend downwards during ribosomal stress but upwards during translation stress. These two gene sets are not daughter specific genes but their time of expression coincides with the time of expression of cell separation genes (http://www.cyclebase.org/).

**Figure 5.**
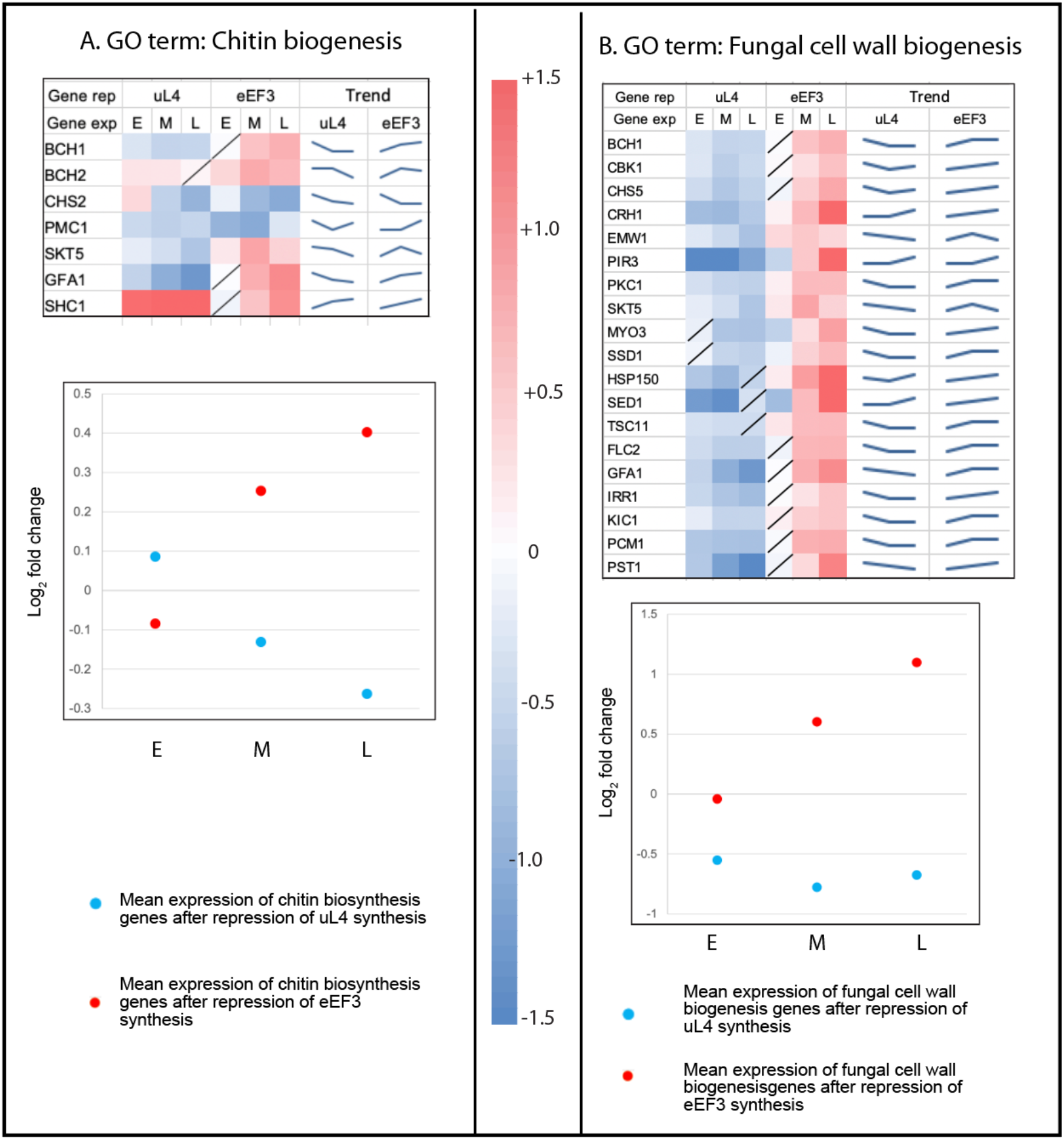
Expression of genes for biogenesis of Chitin and Fungal cell wall. Heatmaps and scatter plots for genes classified under the GO-terms (**A**) Chitin biogenesis, and (**B**) Fungal cell wall biogenesis. For FDR statistics and Trend lines, see legends to Fig 2 and 3.

With few exceptions, the expression of genes in the GO-terms ER to Golgi transport, exocytosis, protein transport, protein transport into the nucleus, and L-amino acid peptidase activity changes less than two-fold up or down, but the trend for all five of these gene sets is positive over the time of translational stress but negative during ribosomal stress (Figs 6 and S5). It may be that the cells under translational stress are compensating for the decreased translational capacity by upregulating the components of the protein delivery system.

**Figure 6.**
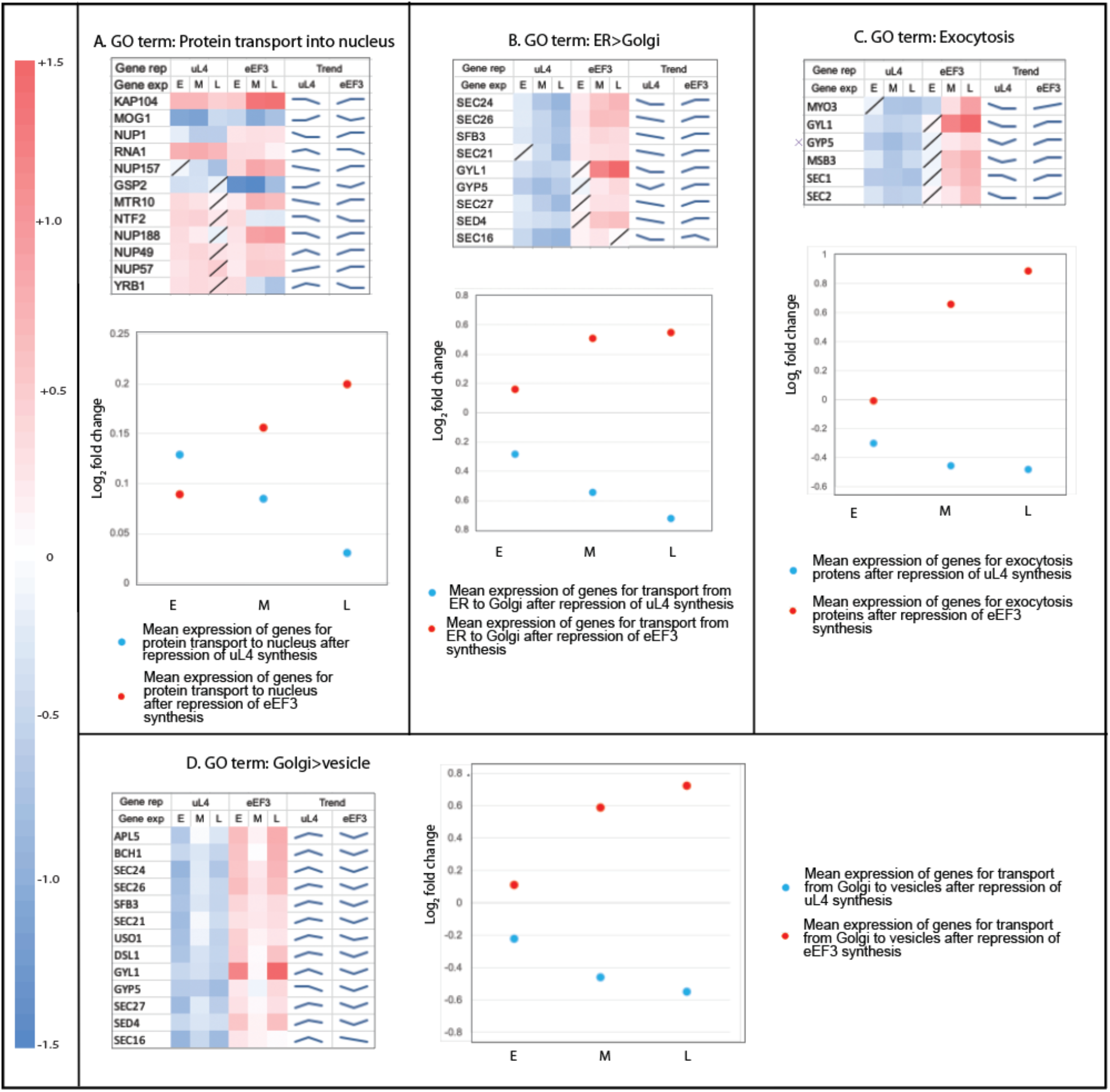
Expression of genes for protein transport. Heatmaps and scatter plots for genes classified under the different protein transport GO-terms. (**A**) Protein transport to the nucleus, (**B**) Protein transport from The Endoplasmic reticulum to the Golgi apparatus, (**C**) Protein transport in Exocytosis, and (**D** Protein transport from Golgi apparatus to vesicles. For FDR statistics and Trend lines, see legends to Fig 2 and 3.

Expression of genes classified under the GO-terms “Response to oxidative stress” and “Endoplasmic reticulum” exhibited increased expression during ribosomal stress and decreased expression during translation stress (Fig 7A-B). *CTA1* and *ERG6* are exceptions and are repressed during ribosomal stress. This may be explained by the fact that the gene products of these two genes are localized to mitochondria.

**Figure 7.**
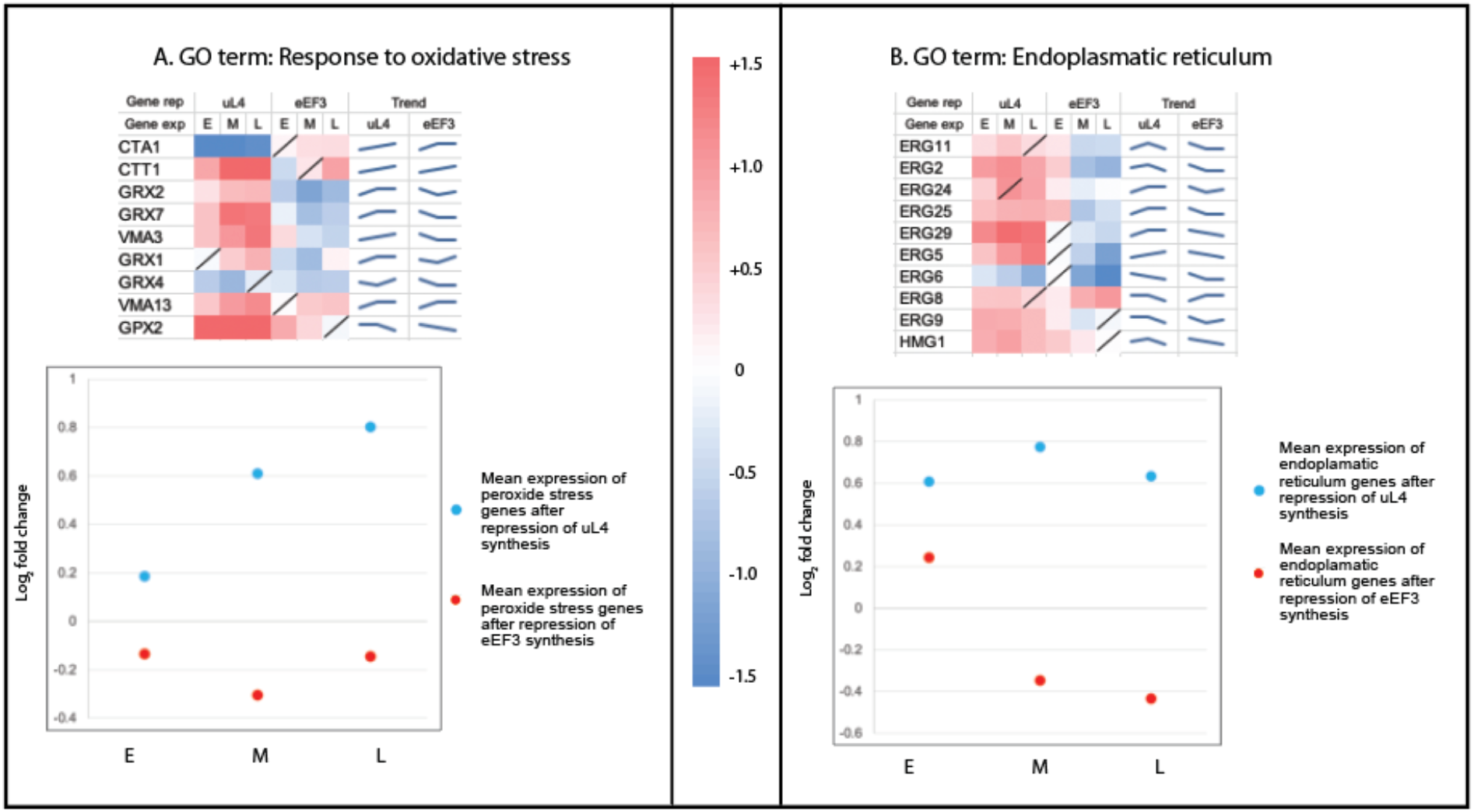
Expression of genes for the formation of the endoplasmic reticulum membrane and response to oxidative stress. Heatmaps and scatter plots for genes under GO-terms associated with various biological processes/molecular functions. (**A**) Endoplasmic reticulum membrane, and (**B**) Response to oxidative stress. For FDR statistics and Trend lines, see legends to Fig 2 and 3.

### GO terms with a similar trend in ribosomal stress and translational stress

#### Membrane turnover

The arrested or delayed cell separation (Shamsuzzaman et al. 2017) suggests that membrane metabolism is changed during both ribosome and translation stress. Related, we noticed that a group of membrane protein genes (*COSxx*) show very similar patterns during both stress forms (Fig 8). The *COS* genes are important for sorting non-ubiquitinated proteins into multivesicular bodies (MBV) for degradation and engulfing of membrane proteins into visible large endosomes (MacDonald et al. 2015), which prompted us to examine vesicles during the abolition of ribosome assembly and ribosome translation. For this purpose, we used strains carrying a GFP-tagged *RAS2*, a marker of plasma membranes. Instead of Pgal-uL4, we used a strain expressing the 60S r-protein eL43 from the *GAL1/10* promoter, which generates the same phenotype as Pgal-uL4 after a shift from galactose to glucose (Gregory et al. 2019). In agreement with the increase in expression of the COS genes during ribosomal and translation stress, we found that the depletion of either eL43 or eEF3 increased the number of endosomes (Fig 9).

**Figure 8.**
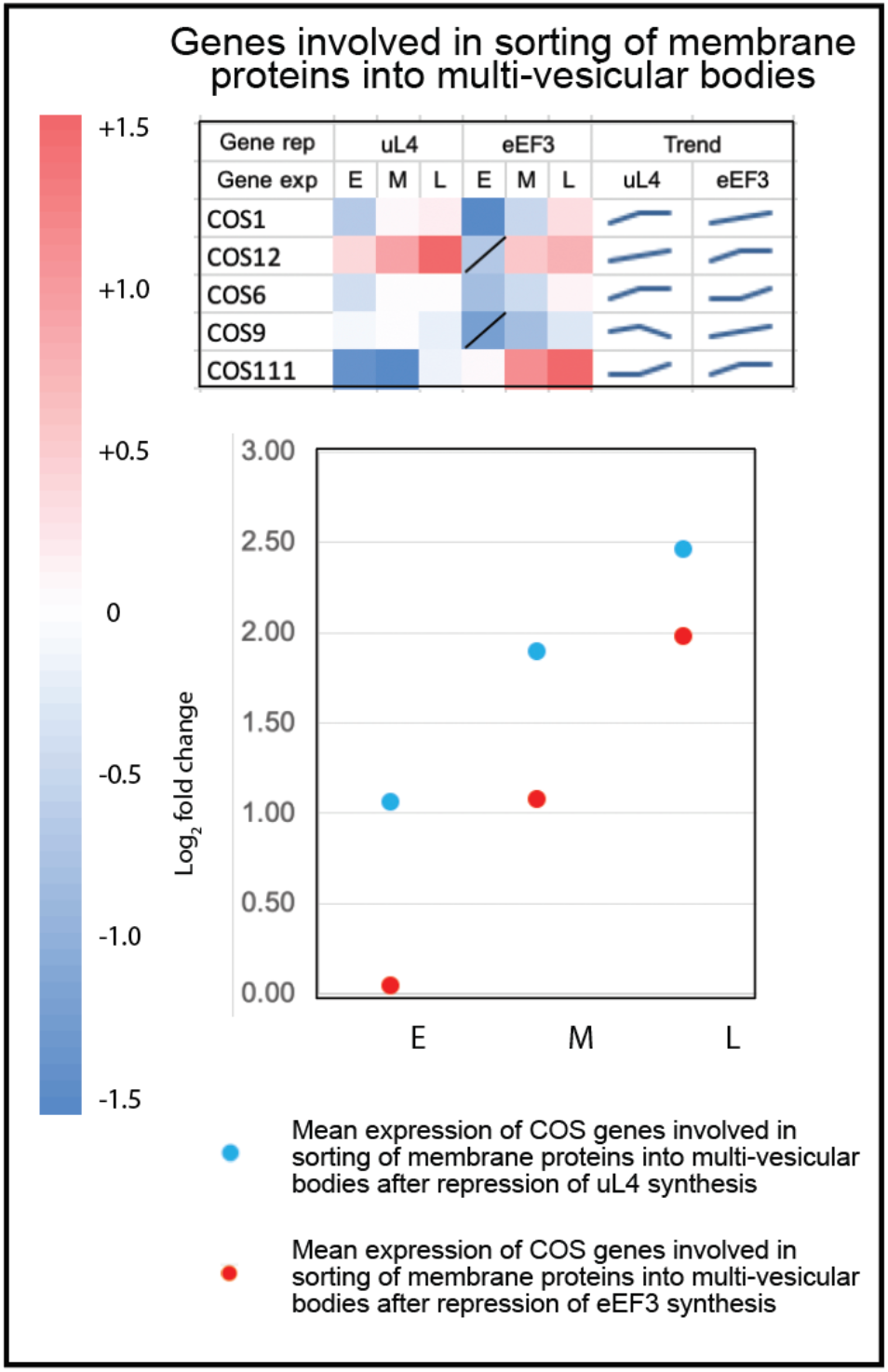
Expression of genes involved in sorting membrane protein into multi-vesicular bodies. Heatmaps and scatterplots for *COS* genes. For FDR statistics and Trend lines, see legends to Fig 2 and 3.

**Figure 9.**
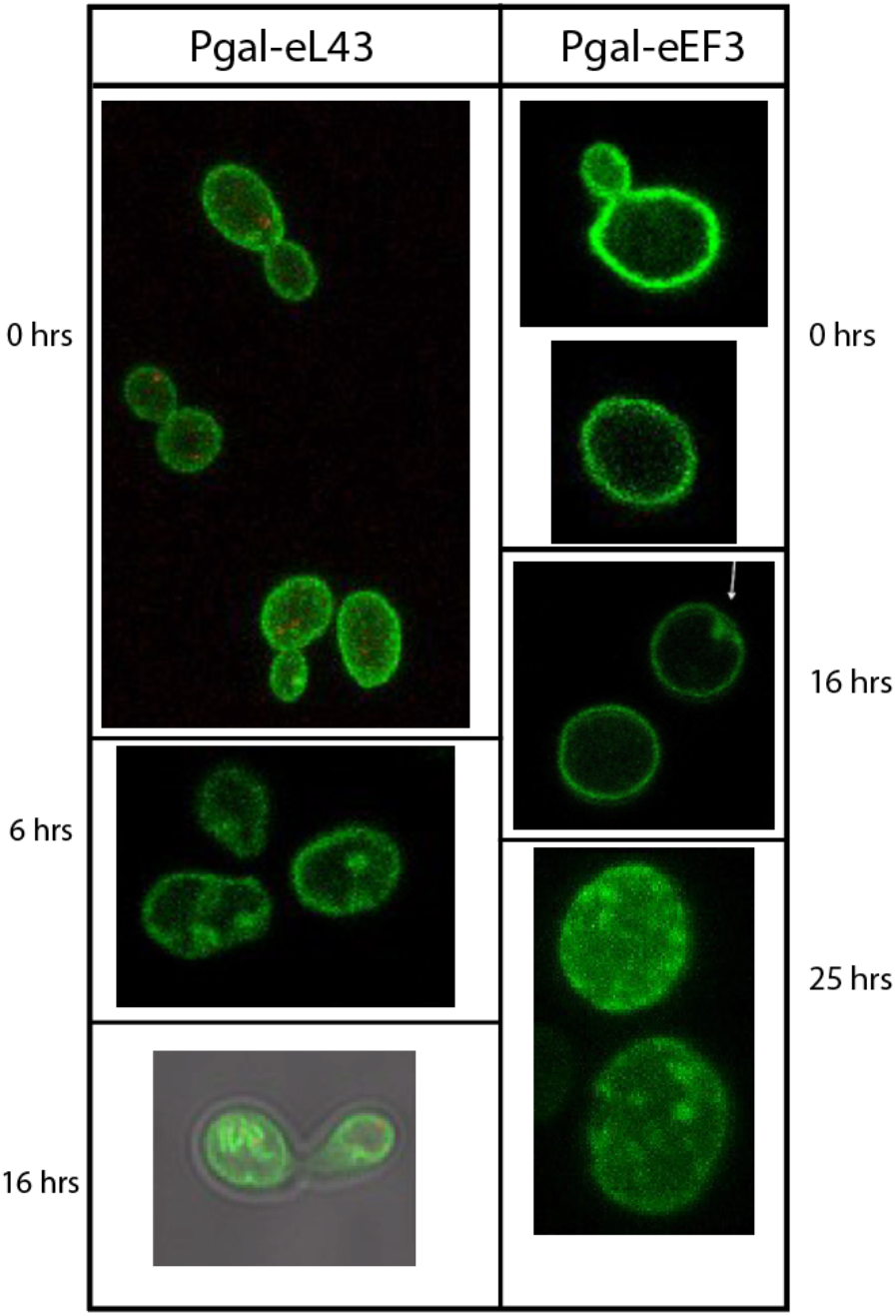
Formation of Multi-vesicular bodies. The membranes were tagged with GFP-Ras2 protein in Pgal-eL43 and Pgal-eEF3. Cells were imaged by confocal microscopy at the indicated times after shifting from galactose to glucose medium.

### Transcription factor overrepresentation analysis of up- and down-regulated DEGs

To identify transcription factors (TF) contributing to the transcriptome changes, we searched for overrepresentation of targets for transcription factors on the lists of DEGs in all samples using the Yeastract database (Monteiro et al. 2008). The top five TFs whose targets are most overrepresented on the lists DEGs of up-regulation and down regulation for a given strain and time point are shown in Table 1 Columns 1 and 5, respectively, and the fraction of genes on each DEG list that are targets for a given TF are shown in Columns 2 and 6. The statistical contribution to gene regulation made by each TF can be deduced by the changes to the fraction of target genes of a given TF the genome that are actually found on the DEG lists (Table 1, Columns 3 and 7). The fraction of genomic target genes for the TF Gat4 found on the DEG list increased over two-fold (9% to 22%) between 2 and 16 hours of ribosomal stress, indicating that Gat4 contribution of differential gene regulation increases with time during ribosomal stress. In contrast, Gat4 targets on the DEG list only increase from about 6 to 9% from the early to late stress stages during inhibition of eEF3 synthesis.

**Table 1.** Analysis of transcription factors.

| Stress type | Sample | Activation |  |  |  | Repression |  |  |  |
| --- | --- | --- | --- | --- | --- | --- | --- | --- | --- |
|  |  | Transcription factor (TF) | #TF target genes/genes on DEG list (Tg/D) | #TF target genes on DEG list/#TF target genes in genome (Td/Tg) | P value | Transcription factor (TF) | #TF target genes/genes on DEG list (Tg/D) | #TF target genes on DEG list/#TF target genes in genome (Td/Tg) | P value |
| Ribosomal | Early | Gat4 | 7.4% | 8.7% | 1.1E-04 | Sut1 | 19.40% | 18.91% | 0.0E+00 |
|  |  | Upc2 | 8.9% | 7.2% | 4.8E-04 | Mig2 | 12.50% | 21.82% | 0.0E+00 |
|  |  | Lys14 | 0.8% | 28.6% | 1.5E-03 | Bas1 | 52.05% | 9.29% | 5.7E-07 |
|  |  | Gzf3 | 7.0% | 6.2% | 8.3E-03 | Rsc30 | 3.92% | 20.59% | 4.8E-06 |
|  |  | Eds1 | 3.9% | 7.4% | 1.0E-02 | Mig1 | 11.01% | 13.02% | 7.6E-06 |
|  | Late | Gat4 | 6.8% | 22.0% | 2.2E-08 | Sut1 | 16.16% | 18.36% | 3.2E-14 |
|  |  | Gis1 | 9.9% | 17.9% | 2.2E-07 | Mig2 | 10.56% | 21.50% | 6.7E-13 |
|  |  | Upc2 | 7.1% | 15.7% | 3.2E-04 | Mot3 | 13.12% | 13.92% | 5.2E-06 |
|  |  | Pdc2 | 1.0% | 26.9% | 2.8E-03 | YPR015C | 4.16% | 18.84% | 5.1E-05 |
|  |  | Msa2 | 0.3% | 50.0% | 3.6E-03 | Mig1 | 10.08% | 13.91% | 6.4E-05 |
| Translation | Early | Gat4 | 17.4% | 5.5% | 1.2E-07 | Bas1 | 66.84% | 8.59% | 0.0E+00 |
|  |  | Mig 2 | 17.4% | 3.9% | 5.9E-06 | Sut1 | 24.61% | 17.27% | 0.0E+00 |
|  |  | Pdc2 | 2.9% | 7.7% | 1.9E-03 | Mig2 | 17.10% | 21.50% | 0.0E+00 |
|  |  | Mot3 | 17.4% | 2.0% | 3.8E-03 | Sok2 | 50.26% | 8.60% | 2.0E-15 |
|  |  | Gis1 | 13.0% | 2.3% | 4.1E-03 | Zap1 | 37.05% | 9.29% | 2.9E-13 |
|  | Late | Rlm1 | 17.9% | 9.4% | 5.3E-07 | Sut1 | 14.48% | 17.09% | 1.9E-10 |
|  |  | Wtm2 | 5.1% | 11.2% | 5.2E-04 | Bas1 | 52.54% | 11.36% | 7.7E-09 |
|  |  | Crz1 | 13.1% | 7.7% | 2.5E-03 | Mig2 | 8.47% | 17.92% | 2.2E-07 |
|  |  | Rgm1 | 8.8% | 8.3% | 3.0E-03 | Mot3 | 12.94% | 14.26% | 6.9E-06 |
|  |  | Gat4 | 5.3% | 9.2% | 5.1E-03 | YPR015C | 4.01% | 18.84% | 9.7E-05 |

Genes targeted by several other TFs are also overrepresented only during one type of stress. The fraction of genomic Upc2 target genes on the DEG list increased from 7% to 16% during ribosomal stress. Upc2 positively regulates ergosterol biosynthesis in response to the cellular level of sterol (Vik A and Rine 2001). Rlm1 is overrepresented during late translation stress. This TF is activated by Pkc1 in the cell wall integrity pathway and positively regulates the expression of cell wall biogenesis genes (Jung et al. 2002). Ergosterol biosynthesis genes are up-regulated in uL4 depletion samples (Fig 7) and cell wall biogenesis genes are upregulated in the eEF3 depletion sample (Fig 5).

Sut1 is acting as a repressor during both types of stress with 15-25% of the genomic target genes found on the DEGs lists throughout the stress periods. This TF positively regulates genes for sterol uptake and negatively regulates gene expression required in filamentous growth in starvation conditions (Foster et al. 2013). We observed elongated buds during ribosomal stress and to a lesser extent during translation stress, indicating filamentous growth ((Thapa et al., 2013) and Md Shamsuzzaman, unpublished). Sut1 repressive activity may inhibit filamentous growth.

Between 18 and 22% of genes targeted by Mig2 as a repressor are found on the DEG lists both early and late during both the stress forms, but this is likely a result of the shift to glucose medium, since Mig2 is known to be involved in glucose repression of the invertase gene *SUC2* (Lutfiyya and Johnston 1996).

## Discussion

We introduced ribosomal and translation stress by repressing the synthesis of r-protein uL4 and Translation Elongation Factor eEF3, respectively. The genes encoding these proteins are repressed by shifting the cultures from galactose to glucose medium causing a shift-up effect (see above), which is stabilized after about an hour (Kief and Warner 1981). Consequently, we have used RNA-seq data in samples collected ≥ 2 hours after the media shift to characterize transcriptomes under stress conditions, and our general findings are summarized in Table 2.

**Table S2.** Summary of the outcome of RNA-seq procedures.

| Gene group | Trend |  | Figure(s) |
| --- | --- | --- | --- |
|  | Ribosomal stress | Translation stress |  |
| Cytoplasmic 60S ribosomal proteins | Up | Down | 3A-B |
| Cytoplasmic 40S ribosomal proteins | Up | Down | 3A-B |
| Ribosome assembly factors (<u>Ribi</u>) | Up | Up | 3B-C |
| Mitochondrial ribosomal proteins | Down | Down | S3 |
| Mitotic Cell cycle | Down | Up | 5A |
| Daughter-specific <u>gluconases</u> | Down | Down | 5C |
| Other daughter-specific genes | Down | Down | S4A |
| Crz1-responsive genes | Up | Up | S4B |
| Chitin biosynthesis genes | Down | Down | 6A |
| Fungal cell wall biogenesis | Down | Up | 6B |
| Protein transport | Down | Up | S5 |
| Protein transport into nucleus | Down | Up | 7A |
| Protein transport ER to Golgi | Down | Up | 7B |
| Protein transport Golgi to vesicles | Down | Up | 7D |
| Exocytosis | Down | Up | 7C |
| Response to oxidative stress | Up | Down | 8A |
| Endoplasmic reticulum membrane | Up | Down | 8B |
| Membrane turnover | Up | Up | 9 |

### Expression of ribosomal protein genes is increased during ribosomal stress and decreased during translation stress

R-protein genes are initially repressed slightly after the imposition of ribosomal stress, but change to a sustained positive trend within 2 hours after cessation of uL4 synthesis (Fig 3A-B(i)). The only major exception is the *RPS22B* gene, one of two paralogous genes encoding r-protein uS8, that is repressed rather than induced during ribosomal stress (Fog 3A) and repressed more rapidly than other r-protein genes during translation stress (Fig S2). The reason for the separate pattern for the *RPS22B* gene may be that this gene has a binding site for an additional transcription factor, Tbf1 (Fermi et al. 2016). The paralogue, *RPS22A*, is upregulated like the remaining r-protein genes (Fig S2). The significance of the differential behavior of the two uS8 genes is not clear, since the encoded proteins only differ in one out of 130 amino acids (I vs L).

R-protein synthesis is repressed after the abolition of rRNA synthesis due to the accumulation of aggregates of r-proteins that fail to be assembled in the absence of de novo rRNA synthesis (Albert et al. 2019a). Most likely, the initial decrease in the mean expression of r-protein genes after cessation of uL4 synthesis is actuated by this mechanism. Ribosomal genes other than the uL4 genes continue to be expressed after shifting Pgal-uL4 to glucose medium (Gregory et al. 2019), but cannot be assembled in the absence of de novo uL4 synthesis, since uL4 is essential for 60S assembly. Unassembled r-proteins are degraded by proteasomes (Gorenstein and Warner 1977; Sung et al. 2016), but the balance between r-protein synthesis and degradation is likely to include a pool of insoluble proteins. Another pathway for repression of r-protein synthesis was revealed by rapamycin inactivation of TORC1, resulting in the release and sequestration of Ifh1 in the CURI complex consisting of the rRNA processing factors Utp22 and Rrp7 rRNA and the CK2 casein kinase (Albert et al. 2016). We do not think that this mechanism is likely to contribute to the initial decreased expression of the r-protein genes since TORC1 should not be inactivated in glucose medium. Whatever the mechanism for the initial decrease in r-protein gene expression, the trend is overruled by a perpetual induction within 2 hours after the imposition of ribosomal stress (Fig 3), which could be due to an increased rate of proteasomal turnover of r-proteins accumulating outside of ribosomal particles (Sung et al. 2016; Rousseau and Bertolotti 2018; Albert et al. 2019a).

It is remarkable that the synthesis of ribosomal components, part of the most resource-demanding process in the cell is upregulated, even though this is futile while ribosome assembly is blocked. Therefore, we did an alternate analysis using DAVID to compare the overrepresentation of differentially expressed genes in our RNA-seq data and in gene expression data collected during heat, oxidative, osmotic, and starvation stress available in yeast genome database (https://spell.yeastgenome.org/search/dataset_listing. As shown in Fig 10 this analysis confirmed the repression of GO-terms related to ribosome formation and other RNA synthesis and processing. In contrast, the same GO-terms were stimulated during ribosomal stress, based on our RNA-seq experiments, thus confirming that ribosome synthesis is stimulated during ribosomal stress, but repressed during environmental stress.

**Figure 10.**
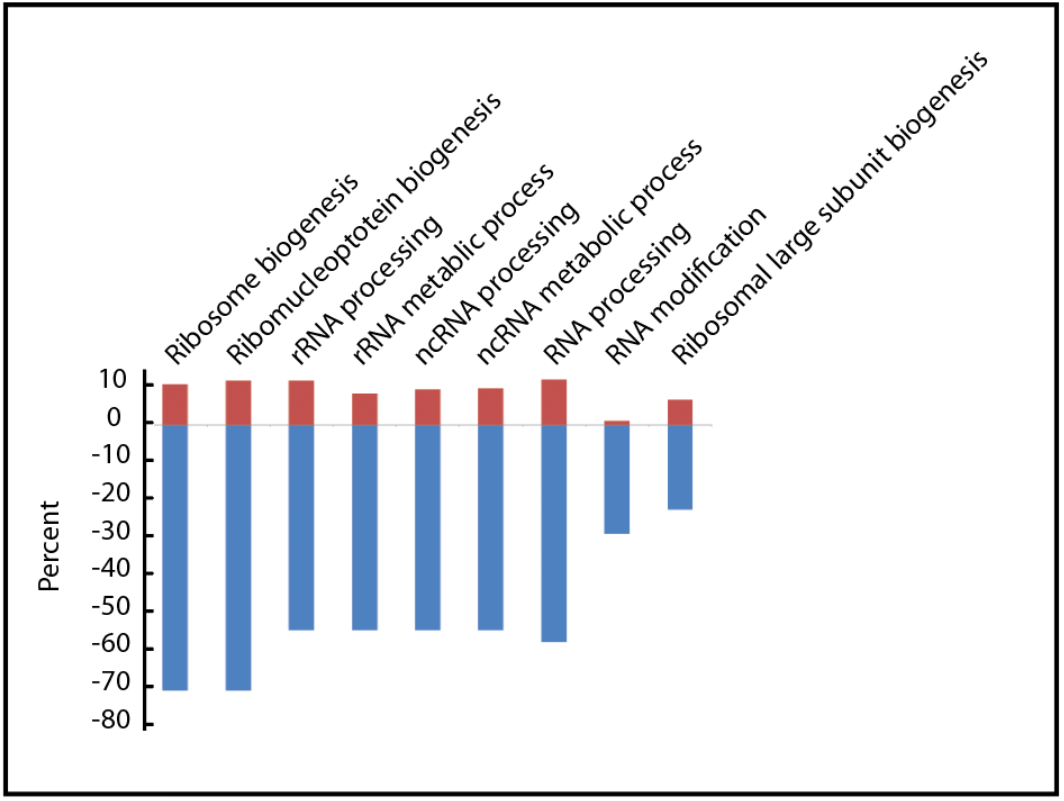
Top GO terms of down-regulated common DEGs in environmental stresses are up-regulated in nucleolar stress. Overrepresentation analysis comparing most common down-regulated DEGs during environmental stress (blue bars) with DEGs in expression data during ribosomal stress (Figs 3–8) shown as red bars. Environmental stress data were obtained from https://spell.yeastgenome.org/search/dataset_listing

During translation stress, the r-proteins initially trend upward, presumably due to the shift-up effect of the shift from galactose to glucose medium. The effect of translation stress can be seen after 2 hours of eEF3 depletion as a monotonous decrease of r-protein gene expression (Fig 3). A likely cause of this is ribosome collisions. When the number of eEF3 molecules falls below the number of ribosomes, an increasing fraction of the ribosomes will become translocation incompetent. Thus, some ribosomes will be immobile on the mRNA while “waiting” for an eEF3 molecule, while others already carry an eEF3 molecule and are translocation competent. As the translocation competent ribosomes catch up to the stationary ribosome the two will collide (Fig 1A(iii)). As the eEF3 concentration continues to decrease, the probability of such collisions increases. Moreover, the fraction of ribosomes in polysomes increases slightly and the cell density continues to increase (Fig. 1B) showing that ribosomes are not stationary on the mRNA, but continues to move and synthesize protein albeit at a decreasing speed (Shamsuzzaman et al. 2017). Thus, eEF3 molecules must rotate randomly between ribosomes, which further increases the probability of ribosome collisions. We, therefore, propose that a likely reason for the progressive repression of r-protein genes during translation stress is an increasing frequency of ribosome collisions with ensuing kinase cascades (Wu et al. 2019; Juszkiewicz et al. 2020; Wu et al. 2020), that ultimately target r-protein transcription. In contrast, it is unlikely that repression emanating from insoluble r-protein aggregates contributes to the downward trend for r-protein genes during translation stress: Since no r-protein gene is specifically repressed, r-proteins are synthesized in a normal coordinated manner synthesis, few, if any, insoluble r-protein aggregates should be formed.

### The Ribi genes are regulated differently than the r-protein genes during both ribosomal and translation stress

The Ribi genes and r-protein genes are part of a large network regulated the Sfp1 transcription factor (Marion et al. 2004; Singh and Tyers 2009; Huber et al. 2011; Bosio et al. 2017; Albert et al. 2019b; Cheng and Brar 2019). However, the Ribi genes are not expressed coordinately with the r-protein genes during either ribosomal or translation stress. In both cases, the Ribi genes show a sustained average increase of about 35%, which eventually fades late in the translation stress response. Importantly, individual Ribi genes are not regulated the same way during ribosomal and translation stress, even though the scatter plots of the **average** regulation are very similar during the two types of stress, excluding the possibility that the regulation of the Ribi genes is simply a result of the change from galactose to glucose medium. Uncoordinated expression of Ribi and r-protein genes was also observed during the proteotoxic response triggered by inhibition of rRNA synthesis discussed above (Albert et al. 2019a). We propose that the large network of Sfp1-controlled genes is composed of gene clusters that can be controlled separately.

We considered the idea that the degree of regulation of individual Rib genes could follow the order in which the assembly factors work in the ribosomal subunit assembly pathways. This might for example be the case if the regulation of genes for factors promoting early steps in each pathway had a gatekeeper role accomplished by tighter regulation of genes for upstream functions, relative to downstream functions in the subunit assembly pathways. However, the regulation of genes for factors catalyzing different steps are not coordinated with the order in which they work in the assembly pathways. Interestingly, the *SDO1* gene, the paralogue of the SBDS responsible for Schwachman-Bodian-Diamond disease, is up-regulated strongly during ribosomal stress. Since Sdo1 is involved in the release of Translation Initiation Factor 6 from precursor 60S in the terminal reaction (Warren 2018) this regulation could be related to retro-regulation mechanisms in rRNA processing reactions (Kofler et al. 2020). It may also be important that Sdo1 appears to have an additional role in the regulation of iron homeostasis (Jain et al. 2020).

### Cells may exit the cell cycle during ribosomal stress

Multiple studies have shown that ribosomal and translation stress arrests or delays the passage through the G1 phase of the cell cycle (Oeffinger and Tollervey 2003; Bernstein and Baserga 2004; Gomez-Herreros et al. 2013; Thapa et al. 2013; Polymenis and Aramayo 2015). Although different steps in the G1 phase could be inhibited dependent on the nature of the interference, the G1 arrest caused under the stress regime used here is caused by inhibition of postmitotic separation of mother and daughter cells (Shamsuzzaman et al. 2017). Dissection of the expression of cell cycle genes strongly suggests that this is due to Crz1-based repression of genes for daughter-specific glucanases and chitinases involved in degrading the cell wall from the daughter side Fig 4.

Despite the similarity in the G1 arrest during ribosomal and translation stress, the overall gene expression patterns suggest that cells may enter a quiescent/stationary mode (sometimes referred to as “G0”) during ribosomal stress, but not during translation stress. Ribosomal stress leads to repression of the genes under the GO-term “Mitotic Cell cycle” (Fig 4) and increased overrepresentation increases for the Gis1 transcription factor (Table 1) that is important for the transition to stationary phase during starvation by inhibiting genes of cell division (Zhang et al. 2009). Moreover, *SHC1*, encoding a chitin synthetase involved in sporulation (Sanz et al. 2002), is heavily increased (Fig 5A).

The gene expression pattern during translation stress is very different and is not compatible with an exit from the cell cycle. This is indicated by the positive trend shown both by genes for the mitotic cell cycle (Fig 4A) and several genes specific to the M/G1 boundary, including genes for cell wall synthesis, show positive trends after depleting eEF3 (Fig 5B).

### Cell wall integrity stress is part of translation stress

Translation stress causes increased expression of many genes for protein transport (Fig 6). We especially point out the continually increased expression of genes for the secretory pathway including ER to Golgi transport, Golgi vesicular transport, exocytosis, and fungal cell wall biogenesis genes, while these genes have a negative trend during ribosomal stress (Fig 6B-D). This pattern of expression of protein transport genes is characteristic of the cell wall integrity pathway (Bulik et al. 2003), suggesting that the cell wall integrity stress is a component of translation stress. The cell integrity pathway activates the SBF transcription complex at the M/G1 boundary of the cell cycle, which in turn induces expression of genes required for G1 to S progression (Kim et al. 2010), and activation of the cell integrity pathway thus harmonizes with the notion that cells are not in a quiescent mode during translation stress as they appear to be during ribosomal stress.

The decreased expression of the protein transport genes during ribosomal stress is compatible with the view that ribosome formation is required for protein secretion. Previous results from the Warner lab showed that mutations interrupting the secretory pathway repress both r-protein and rRNA genes (Mizuta and Warner 1994; Nierras and Warner 1999). Thus, it appears that ribosome biogenesis and protein secretion and mutually dependent on each other. The coupling of ribosome formation to protein secretion depends on the Pkc1 kinase, which is required for remodeling of cell walls (Nierras and Warner 1999; Li et al. 2000). Genes under the cell wall GO-term are induced during translation stress but repressed during ribosomal stress (Fig 5B), which is compatible with the overrepresentation of targets for the Rlm1 transcription factor after 26 hours of eEF3 depletion (Table 1) because Rim1 works as an effector of the Pkc1 signaling pathway and turns on expression of cell wall biogenesis genes (Jung et al. 2002). Together, our results are compatible with the notion that the cell integrity pathway is activated during translational stress via a mechanism involving Pkc1.

### Oxidative stress and acidification of the cell during ribosomal stress

Several genes under the GO-term “Response to oxidative stress” are increased sharply during ribosomal stress, suggesting that oxidative stress is a component of ribosomal stress (Fig 7A). This includes a continual positive trend of *CTT1*, encoding for a cytoplasmic catalase that neutralizes small peroxides like H2O2. In contrast, the gene for the mitochondrial catalase *CTA1* is repressed. Also upregulated are the genes for Grx and Gpx enzymes that neutralize large molecules hydroperoxide like tert-butyl hydroperoxide (t-BHP) and lipid hydroperoxide (Avery and Avery 2001), including a 7-fold induction of the gene for Gpx2 phospholipid hydroperoxide glutathione peroxidase as early as in 2h (Fig 7A). The increased expression of lipid peroxidases suggests that the cell membrane components are oxidized during ribosomal stress, which is also indicated by the increased expression of *GEX1* whose product functions in the export of excess oxidized glutathione and import of H^+^ (Dhaoui et al. 2011). Further support for oxidative stress during ribosome assembly defects comes from the increased expression of VMA3 and VMA13 that both are involved in vacuolar acidifications and are induced under oxidative stress (Fig 7A) (Belli et al. 2004).

Evidence of defects in plasma membrane formation also comes from the increased expression of the *ERG* genes (Fig 7B) and enrichment of the Upc2 ergosterol sensor (Table 1) that function in ergosterol biosynthesis required for membrane formation and the function of the vacuolar ATPase (Smith et al. 1996; Zhang et al. 2010). Together, these observations suggest that cells under ribosomal stress may have an ergosterol deficiency causing deficient vacuolar acidification leading to increased cellular pH level and reactive oxygen species (ROS) level (Niles and Powers 2014).

### Genes that trend similarly during both types of stress

#### Elevated expression of genes involved in the degradation of the membrane during both of these stress conditions

In contrast to the many examples of differential gene expression during ribosomal and translation stress, the 12 *COS* genes are upregulated in cells depleted for either uL4 or eEF3 (Fig 8). This gene set is induced during nutrient stress, causing down-regulation of cell surface proteins, predominantly GIP anchored membrane proteins (MacDonald et al. 2015). Cos proteins work as trans-ubiquitination signals for sorting of membrane proteins into multivesicular bodies for degradation and their expression leads to the formation of large endosomal vesicles (MacDonald et al. 2015). Using fluorescence microscopy of cells expressing GFP-tagged Ras2 membrane protein, we observed similar large endosomal vesicles in cells depleted for either eL43 (which has the same effect on ribosome assembly as uL4 depletion (Gregory et al. 2019)) or eEF3 (Fig 9), presumably a result of the increase expression of Cos proteins. We speculate that both ribosome and translation stress membrane proteins are down-regulated from the cell surface including TORC2 and its mediators Rom2, Slm1, and Slm2. That in turn, may lead to downregulation of TORC2-Ypk1/Ypk2 signaling pathway causing actin depolarization and elevation of ROS (Niles and Powers 2012). Together the trends of genes for detoxification of peroxides and vacuole acidification suggest membrane proteins are oxidized and need to be replaced frequently.

## Concluding Comments

We have compared the transcriptomes developing after inducing two different types of stress that both reduce translation capacity, but by different mechanisms. In the “translating stress” mode the Translation Elongation Factor eEF3 is depleted, which distorts ribosome function without affecting the number of ribosomes. In the “ribosomal stress” mode, the synthesis of an r-protein is stopped, leading to a decrease in ribosome numbers without affecting the functionality of the remaining ribosomes.

Comparing the responses to the two assaults, summarized in Table 2, leads to three major conclusions: (i) There is not a universal stress response to decreased protein synthesis capacity; (ii) the changes in transcriptome are not static, but evolve with time; (iii) a separate mosaic of expression of genes under different GO-terms develops in response to each of the two stress forms. That is, the stress response and cell fate are not determined by the translation capacity per se, but by the **process** that lowers the translation capacity.

We suggest that mutations in ribosomal genes are likely to simultaneously affect the synthesis of ribosomes **and** the ribosome acuity for a normal, accident-free translation process. This, in turn, generates unique combinations of ribosomal and translation stress for each individual mutation in ribosomal genes and results in a specific transcriptome for each mutation. This may be relevant to the development of disease patterns in individual ribosomopathies.

## Materials and Methods

### Strains and growth conditions

All yeast strains used were derived from BY4741 (Brachmann et al. 1998). In each strain, the gene for r-protein uL4 or eL43, or Translation Elongation Factor 3 (eEF3 encoded by *YEF3*) was under the control of the *GAL1/*10 promoter (Table S1). We refer to these strains as Pgal-uL4, −eL43, and −eEF3, respectively. The construction of a derivative of Pgal-eL43 carrying a GFP-tagged RAS2 gene has been described previously (Shamsuzzaman et al. 2017).

Cultures were grown asynchronously in 1% yeast extract, 2% peptone, 2% galactose (YEPGal) at 30°C until mid–log phase (OD_600_=0.8–1.0 corresponding to 1.5–2 × 10^7^ cells/ml) and were then shifted to glucose medium (Shamsuzzaman et al. 2017). All experiments were done in biological triplicates. Cultures were diluted as necessary with pre-warmed media to keep the OD600 <1.0. We did not expose the cultures to cycloheximide prior to harvest which can influence both polysome content and mRNA abundance (Helser et al. 1981; Cheng and Brar 2019).

### RNA preparation and analysis

Samples (1 OD unit) were taken from cultures before and at the indicated times after the shift to glucose medium. Total RNA was extracted using the Ribopure Yeast kit (ThermoFisher, USA) following the manufacturer’s protocol (https://tools.thermofisher.com/content/sfs/manuals/1926ME.pdf). RNA integrity was checked by Bio-Analyzer (Agilent Technologies, USA). One of the three Pgal-eEF3 12h samples did not pass quality control and was not sequenced. RNA samples were enriched for mRNA using poly-A attached magnetic beads followed by Paired-end RNA-seq library preparation using the TruSeq RNA Sample Prep kit (Illumina). One hundred nucleotides were sequenced using the HiSeq platform (Illumina) from both ends of each cDNA and sequence reads were aligned against the *Saccharomyces cerevisiae* S288c reference genome using Tophat2 aligner (Kim et al. 2013). Sequence annotation and read counts per gene were generated using HTseq (Anders et al. 2015) and Subread (Liao et al. 2013) and compared for consistency. We obtained a mean of 2.3-3.1*10^7^ and 2.5-3.3*10^7^ reads mapping to the *S. cerevisiae* BY4741 genome for each sample of Pgal-uL4 and Pgal-eEF3, respectively (Table S2). DESeq2 (Love et al. 2014) was used for calculating differential gene expression between RNA samples relative to the 0-hour sample. A gene was classified as a differentially expressed gene (DEG) if the false discovery rate (FDR) value was <0.05 and the absolute log_2_ (fold change) value was ≥1. The Python Sci-kit learn was used for Principal Component Analysis. Heat maps for figures in the text were generated by Plateau software, while the heat maps in the Supporting Materials was generated by Excel and Python Seaborn library. Heatmaps and scatterplots in the text show genes for which ≥5 of the 6 time points had an FDR ≤0.05. In the Supporting Materials heatmaps show all genes in the indicated GO-term and the scatterplots compare the average LFC for genes with ≥5 of 6 timepoints have FDR≤0.05 with the average LFC for genes with FDR≤0.05 at <5 time points.

### Gene ontology enrichment and overrepresentation analysis

Overrepresentation analysis was performed by mapping differentially expressed genes (DEGs) in stress and control samples to GO terms using geneontology.org (http://www.geneontology.org/; validation date 04/28/2017 and gaf-version 2.0) or to pathway terms available in KEGG, Reactome, Wikipathways, and YeastCyc using the interface of ConsensusPathDB (Kamburov et al. 2011). The number of DEGs mapped to each term were counted and compared to DEGs for all genes of yeast genome. A hypergeometric test was performed and the terms with adjusted p value / q value / False discovery rate < 0.05 were defined as over-represented terms. For enrichment analysis, all genes detected by RNA-seq were ranked by the log_2_ of fold change value (LFC) between the stress and control samples and mapped to GO terms. Wilcoxon signed-rank test was performed, and probabilities were calculated to determine if the combined LFC between genes in a functional group were by chance. The terms with adjusted p-value, q-value, and false discovery rate < 0.05 were defined as overenriched terms. Overrepresentation analysis in Fig 10 was done as described (Le Cao et al. 2014) https://www.bioconductor.org/packages/release/bioc/vignettes/RankProd/inst/doc/RankProd.pdf.

### Identification of gene expressed most differently during ribosomal and translation stress

To identify the genes that were regulated most differently during ribosomal and translational stress we excluded the genes with <50 reads and ranked the rest in ascending order of fold change of expression for each time point, subtracted the gene rank values for individual genes in the Pgal-uL4 samples from the rank values of the Pgal-eEF3 the corresponding sample, and calculated the calculate rank value difference (RVD) for all three time points. The absolute values and actual values of RVDs of each gene were summed over three time points to calculate the absolute rank-sum and rank-sum, respectively. Finally, the absolute rank-sum was subtracted from absolute value of rank-sum to calculate the rank sign. Genes with the highest absolute rank-sum values differ the most in the fold change between the two strains (Pgal uL4 and - eEF3) over the three time points of treatment. The genes with a high rank sign values are the genes which had higher rank in uL4 relative to Tef3 at a specific time point, but changes to lower rank relative to Tef3 at other time points and vice versa. The genes were ranked according to the absolute rank-sum of differences between uL4 and Tef3 depletion samples the generate a list of top differentiating genes.

### Transcription factor (TF) overrepresentation analysis

The Yeastract database was used for TF overrepresentation analysis (Monteiro et al. 2008).

**Confocal microscopy was performed as** previously described (Shamsuzzaman et al. 2017). Whole-cell images of vesicles were obtained by projecting Z-stack images onto a single plane (“collapsing the Z-stack”).

## Acknowledgments

This work was supported by 0920578 grant from the National Science Foundation to Janice M Zengel and LL. Additional funding was provided by an internal appropriation from the University of Maryland, Baltimore County to LL (no grant number). We thank Benedikte Traasdahl for help with the manuscript.

---

**Figure S1.**
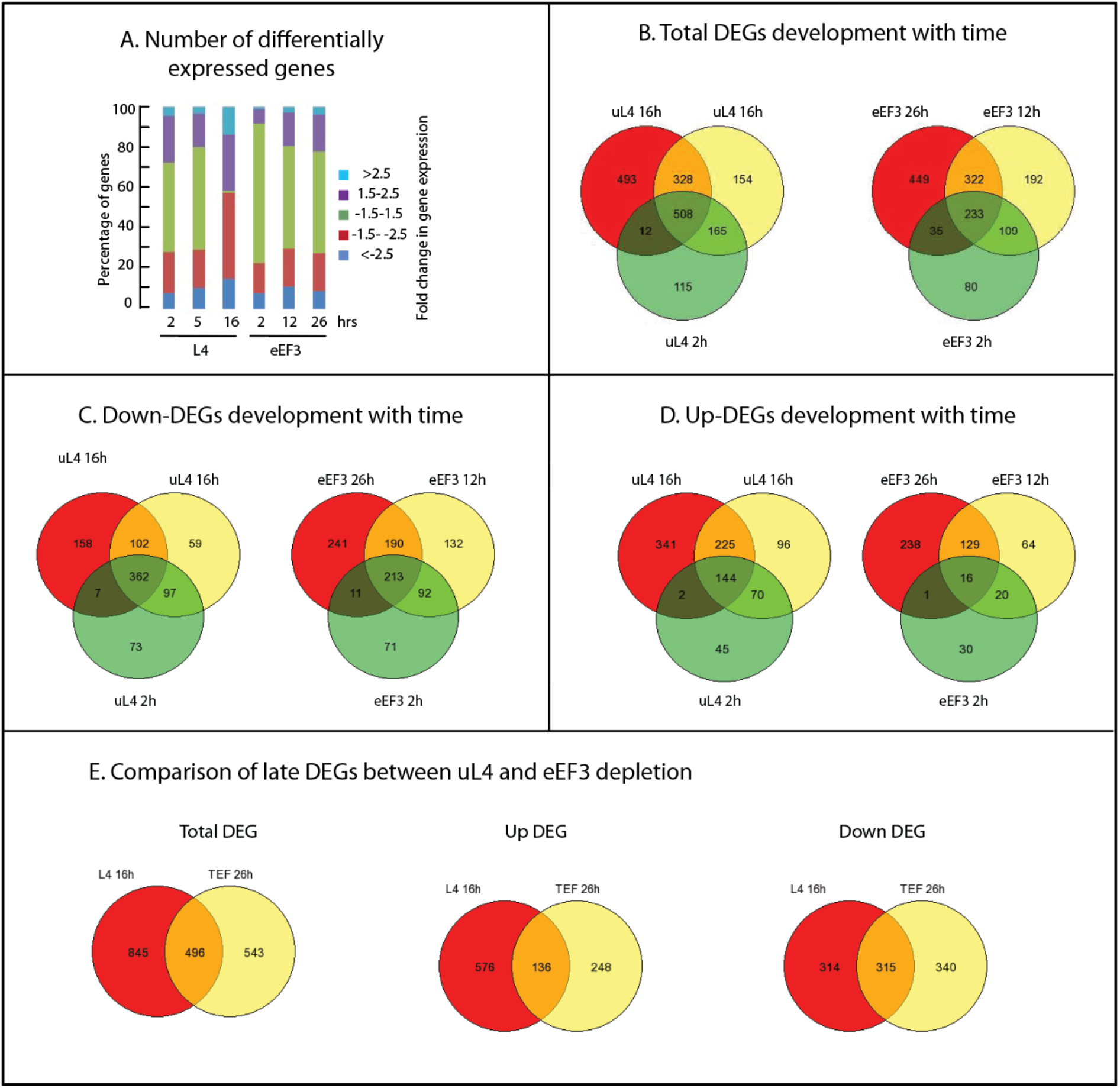
General trends in gene regulation during ribosomal and translation stress. (**A**) Bar plots showing the distribution of fold changes in gene expression after repressing the uL4 or eEF3 synthesis for the indicated times. (**B-D**) Venn diagram comparing unique DEGs in samples collected at different times. (**E**) Comparison of unique DEGs in late samples from Pgal-uL4 and Pgal-eEF3.

**Figure S2.**
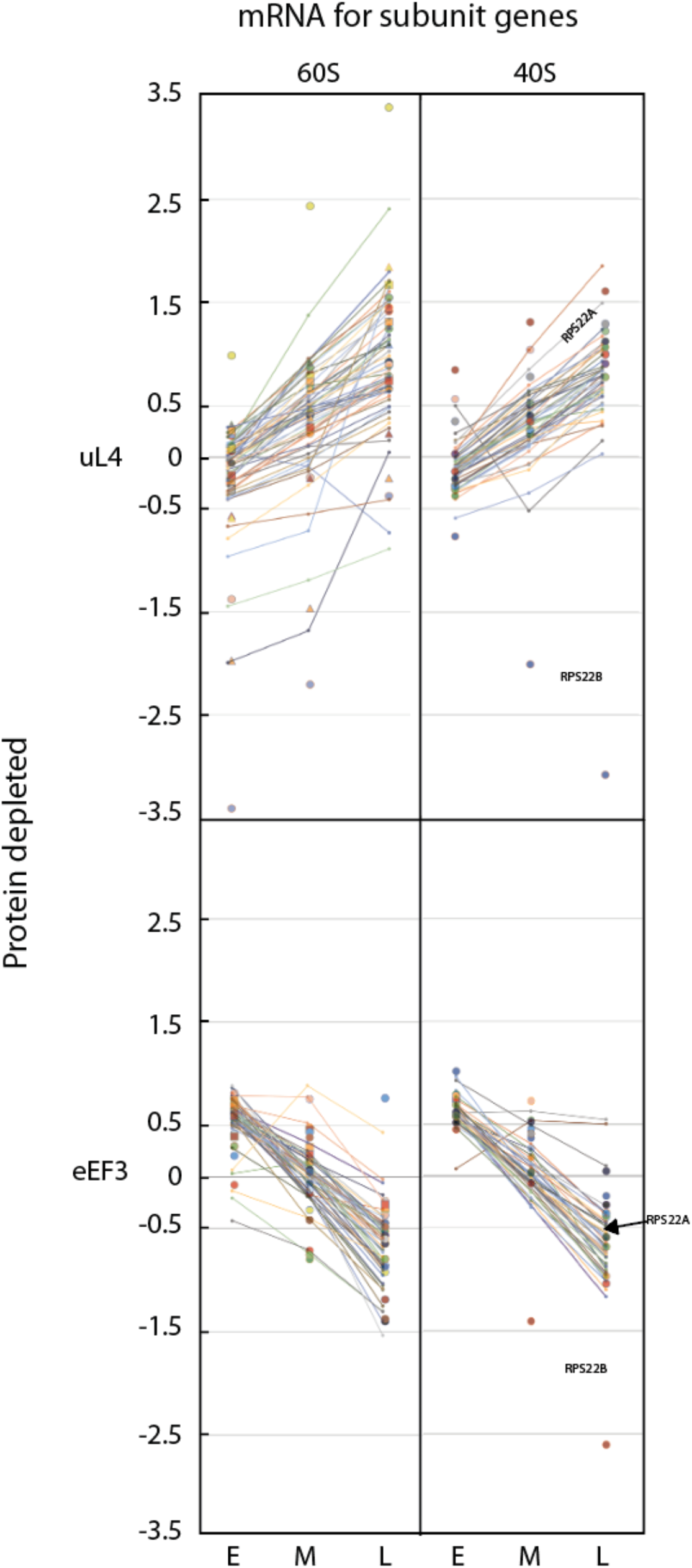
Comparison of the expression of r-protein genes with FDR≤0.05 and >0.05. Log_2_ fold change for genes with FDR≤0.05 are shown as circles and triangles. Genes with FDR>0.05 are shown as lines. The data for RPS22B are identified (see text)

**Figure S3.**
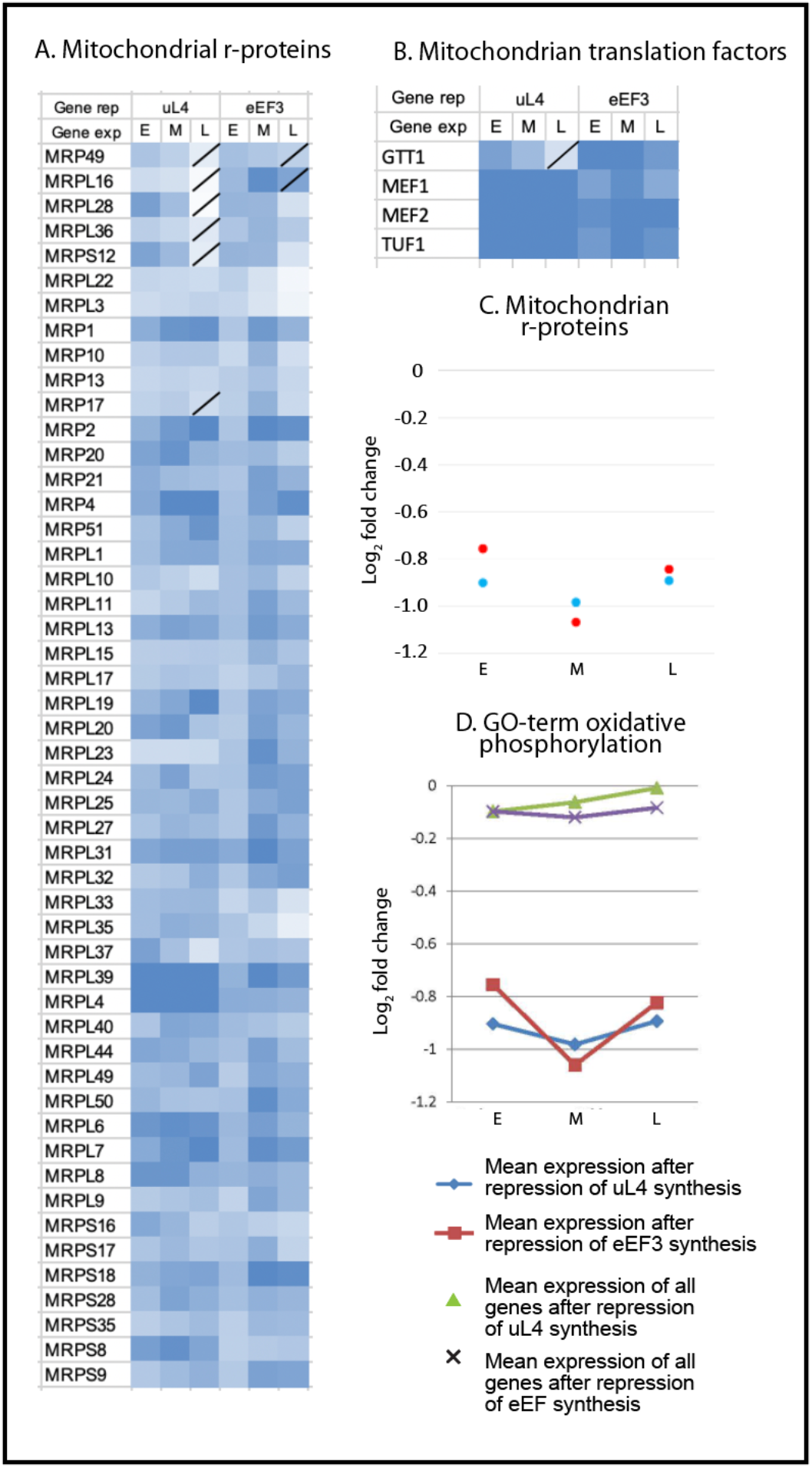
Expression of genes for mitochondrial translation and oxidative phosphorylation. (**A**) Heatmap for expression of mitochondrial ribosomal protein genes. (**B**) Heatmap for expression of mitochondrial translation factors. (**C**) Scatterplots for expression of mitochondrial ribosomal protein genes and (**D**) Heatmap for genes under the GO-term “Oxidative phosphorylation”. For FDR statistics and Trend lines, see legend to Fig 2 and 3.

**Figure S4.**
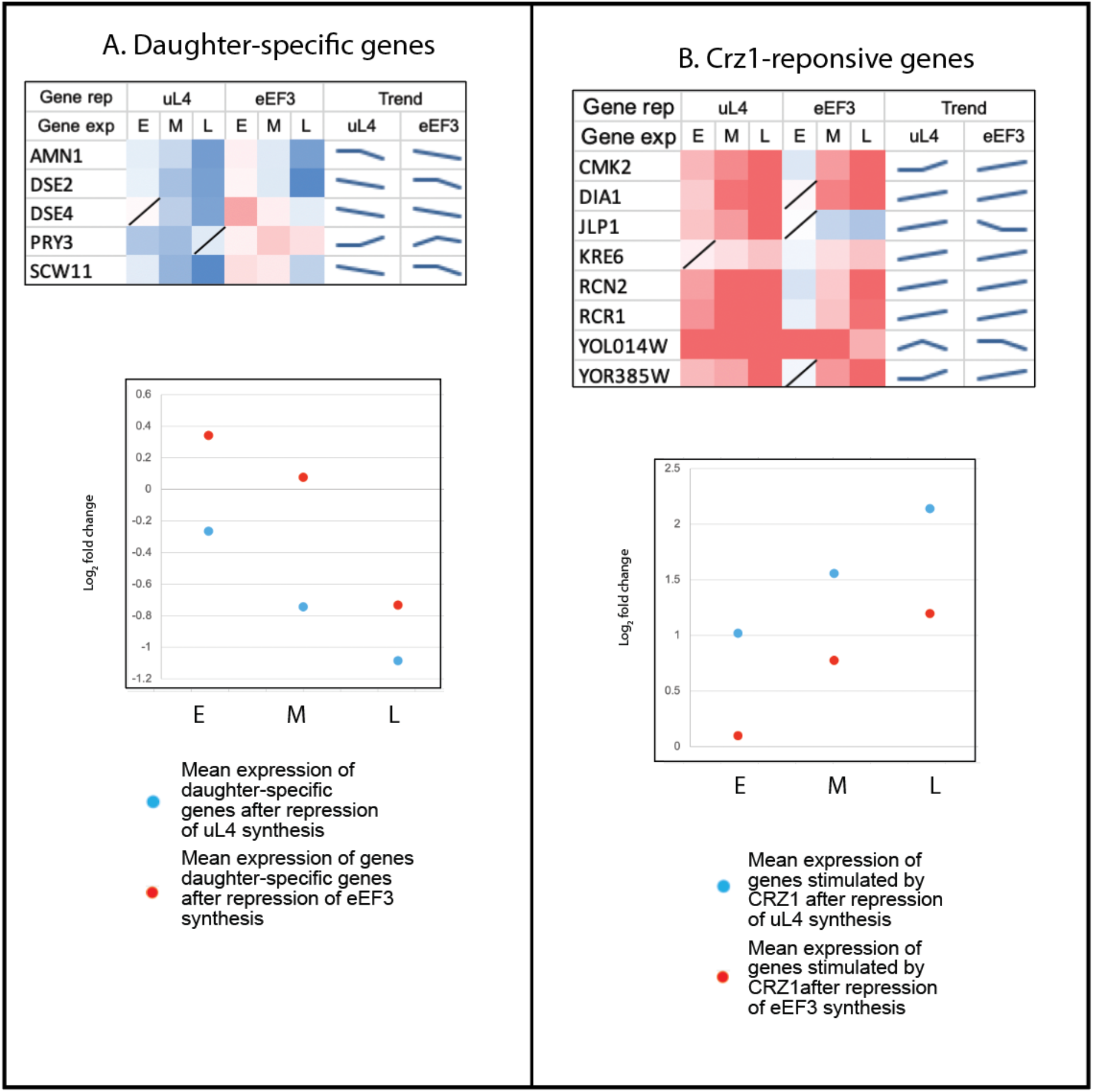
Expression of daughter genes and Crz1-responsive genes. Heatmaps and scatter plots for (**A**) expression of daughter-specific genes and (**B**) genes regulated by the Crz1 transcription factor. For FDR statistics and Trend lines, see legend to Fig 2 and 3.

**Figure S5.**
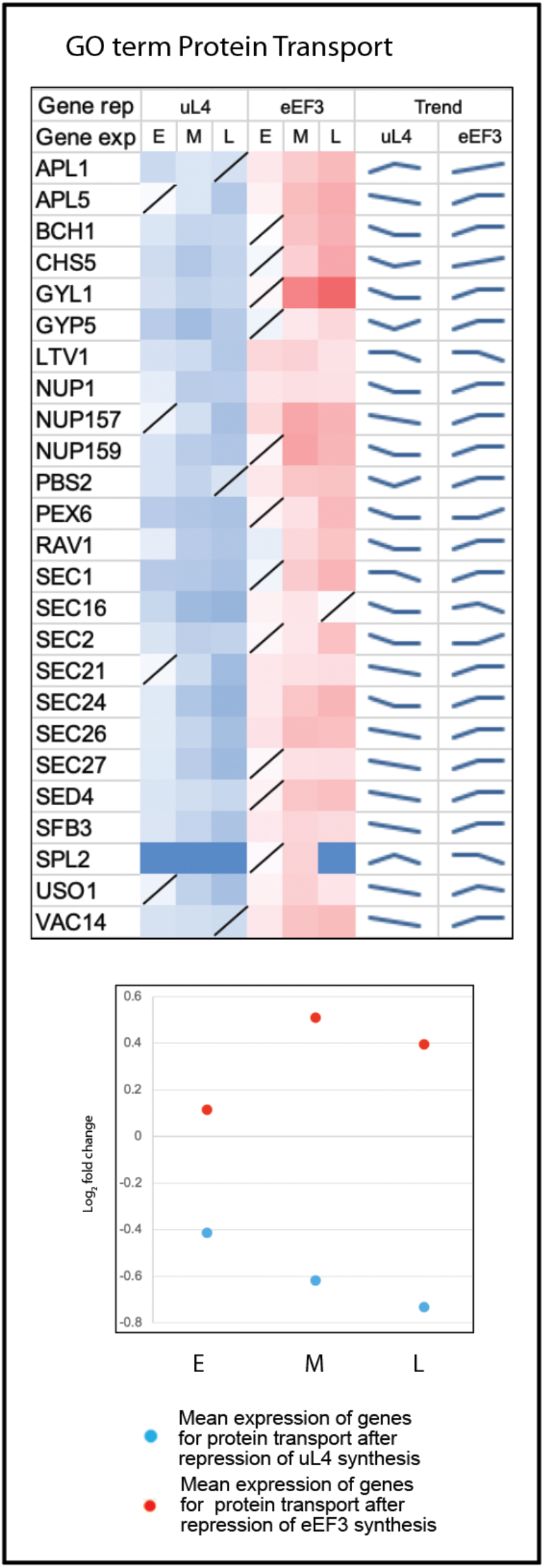
Expression of protein transport genes. Heatmap and scatter plot for the expression of genes cataloged under GO-term “protein transport”.

**Table S1.** Strains used.

| Strain name | Protein encoded by gal-regulated gene | Reference |
| --- | --- | --- |
| Pgal-uL4 | Ribosomal protein uL4/L4 | (Pöll et al. 2009) |
| Pgal-eEF3 | Translation Elongation Factor eEF3 | (Shamsuzzaman et al. 2017) |
| Pgal-eL43 | Ribosomal protein eL43/L43 | (Shamsuzzaman et al. 2017) |

**Table S2.**
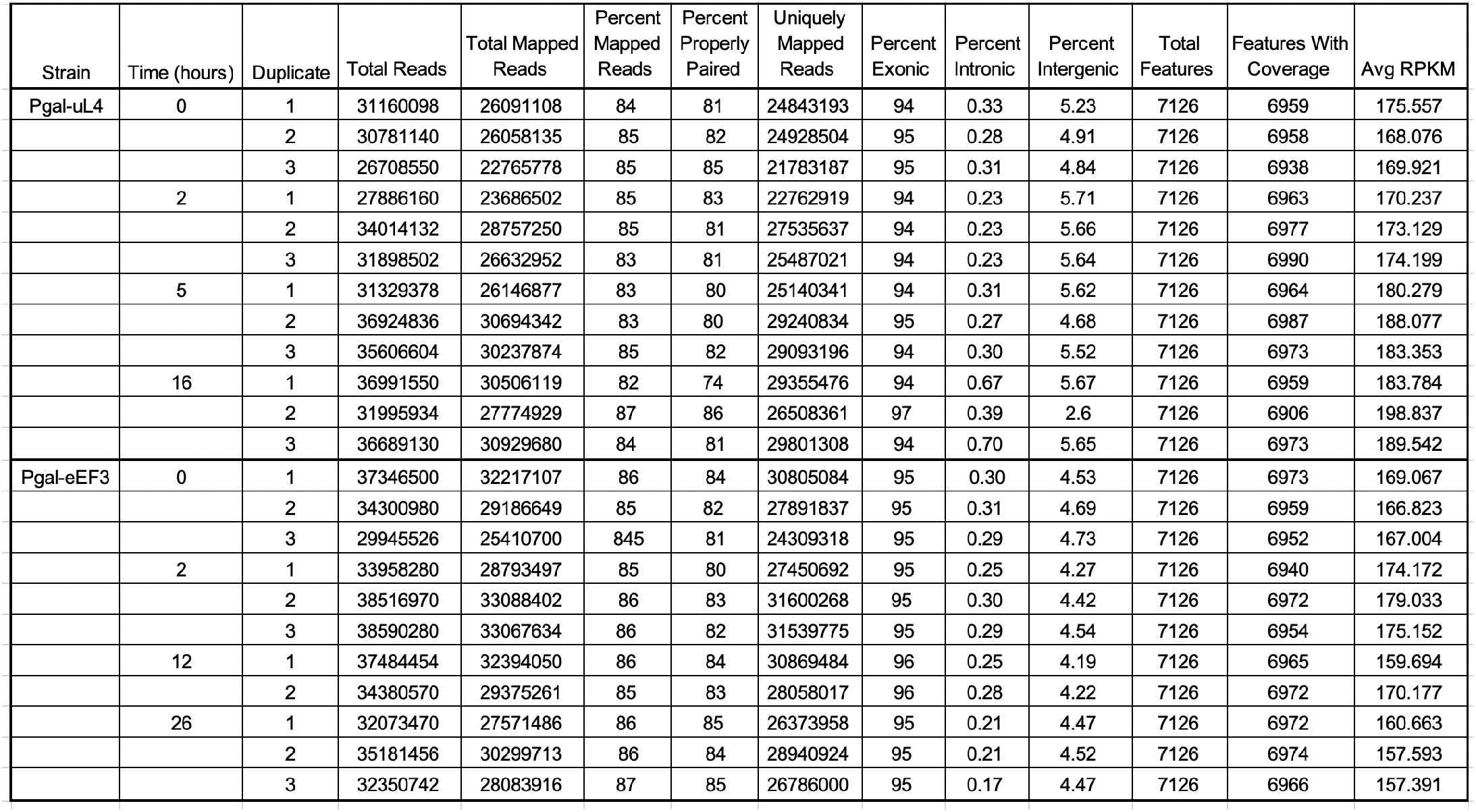
Summary of the outcome of RNA-seq procedures.

## References

Albert B, Knight B, Merwin J, Martin V, Ottoz D, Gloor Y, Bruzzone MJ, Rudner A, Shore D. 2016. A Molecular Titration System Coordinates Ribosomal Protein Gene Transcription with Ribosomal RNA Synthesis. Mol Cell 64: 720–733.

Albert B, Kos-Braun IC, Henras AK, Dez C, Rueda MP, Zhang X, Gadal O, Kos M, Shore D. 2019a. A ribosome assembly stress response regulates transcription to maintain proteome homeostasis. Elife 8.

Albert B, Tomassetti S, Gloor Y, Dilg D, Mattarocci S, Kubik S, Hafner L, Shore D. 2019b. Sfp1 regulates transcriptional networks driving cell growth and division through multiple promoter-binding modes. Genes Dev 33: 288–293.

Anders S, Pyl PT, Huber W. 2015. HTSeq--a Python framework to work with high-throughput sequencing data. Bioinformatics 31: 166–169.

Armistead J, Triggs-Raine B. 2014. Diverse diseases from a ubiquitous process: the ribosomopathy paradox. FEBS Lett 588: 1491–1500.

Aubert M, O’Donohue MF, Lebaron S, Gleizes PE. 2018. Pre-Ribosomal RNA Processing in Human Cells: From Mechanisms to Congenital Diseases. Biomolecules 8.

Avery AM, Avery SV. 2001. Saccharomyces cerevisiae expresses three phospholipid hydroperoxide glutathione peroxidases. J Biol Chem 276: 33730–33735.

Ban N, Beckmann R, Cate JH, Dinman JD, Dragon F, Ellis SR, Lafontaine DL, Lindahl L, Liljas A, Lipton JM et al. 2014. A new system for naming ribosomal proteins. Current opinion in structural biology 24: 165–169.

Bassler J, Hurt E. 2019. Eukaryotic Ribosome Assembly. Annu Rev Biochem 88: 8.1–8.26.

Belli G, Molina MM, Garcia-Martinez J, Perez-Ortin JE, Herrero E. 2004. Saccharomyces cerevisiae glutaredoxin 5-deficient cells subjected to continuous oxidizing conditions are affected in the expression of specific sets of genes. J Biol Chem 279: 12386–12395.

Bernstein KA, Baserga SJ. 2004. The small subunit processome is required for cell cycle progression at G1. Mol Biol Cell 15: 5038–5046.

Bohnsack KE, Bohnsack MT. 2019. Uncovering the assembly pathway of human ribosomes and its emerging links to disease. EMBO J 38: e100278.

Bosio MC, Fermi B, Dieci G. 2017. Transcriptional control of yeast ribosome biogenesis: A multifaceted role for general regulatory factors. Transcription 8: 254–260.

Brachmann CB, Davies A, Cost GJ, Caputo E, Li J, Hieter P, Boeke JD. 1998. Designer deletion strains derived from *Saccharomyces cerevisiae* S288C: a useful set of strains and plasmids for PCR-mediated gene disruption and other applications. Yeast 14: 115–132.

Bulik DA, Olczak M, Lucero HA, Osmond BC, Robbins PW, Specht CA. 2003. Chitin synthesis in Saccharomyces cerevisiae in response to supplementation of growth medium with glucosamine and cell wall stress. Eukaryotic cell 2: 886–900.

Bursac S, Brdovcak MC, Donati G, Volarevic S. 2014. Activation of the tumor suppressor p53 upon impairment of ribosome biogenesis. Biochim Biophys Acta 1842: 817–830.

Cheng Z, Brar GA. 2019. Global translation inhibition yields condition-dependent de-repression of ribosome biogenesis mRNAs. Nucleic Acids Res 47: 5061–5073.

Colman-Lerner A, Chin TE, Brent R. 2001. Yeast Cbk1 and Mob2 activate daughter-specific genetic programs to induce asymmetric cell fates. Cell 107: 739–750.

Costanzo MC, Fox TD. 1990. Control of mitochondrial gene expression in Saccharomyces cerevisiae. Annu Rev Genet 24: 91–113.

Deisenroth C, Zhang Y. 2011. The Ribosomal Protein-Mdm2-p53 Pathway and Energy Metabolism: Bridging the Gap between Feast and Famine. Genes Cancer 2: 392–403.

Dhaoui M, Auchere F, Blaiseau PL, Lesuisse E, Landoulsi A, Camadro JM, Haguenauer-Tsapis R, Belgareh-Touze N. 2011. Gex1 is a yeast glutathione exchanger that interferes with pH and redox homeostasis. Mol Biol Cell 22: 2054–2067.

Fancello L, Kampen KR, Hofman IJ, Verbeeck J, De Keersmaecker K. 2017. The ribosomal protein gene RPL5 is a haploinsufficient tumor suppressor in multiple cancer types. Oncotarget 8: 14462–14478.

Farley-Barnes KI, Ogawa LM, Baserga SJ. 2019. Ribosomopathies: Old Concepts, New Controversies. Trends Genet 35: 754–767.

Fermi B, Bosio MC, Dieci G. 2016. Promoter architecture and transcriptional regulation of Abf1-dependent ribosomal protein genes in Saccharomyces cerevisiae. Nucleic Acids Res 44: 6113–6126.

Farley-Barnes KI, Ogawa LM, Baserga SJ. 2017. Multiple roles of the general regulatory factor Abf1 in yeast ribosome biogenesis. Current genetics 63: 65–68.

Foster HA, Cui M, Naveenathayalan A, Unden H, Schwanbeck R, Höfken T. 2013. The zinc cluster protein Sut1 contributes to filamentation in Saccharomyces cerevisiae. Eukaryot Cell 12: 244–253.

Gasch AP, Spellman PT, Kao CM, Carmel-Harel O, Eisen MB, Storz G, Botstein D, Brown PO. 2000. Genomic expression programs in the response of yeast cells to environmental changes. Mol Biol Cell 11: 4241–4257.

Gomez-Herreros F, Rodriguez-Galan O, Morillo-Huesca M, Maya D, Arista-Romero M, de la Cruz J, Chavez S, Munoz-Centeno MC. 2013. Balanced production of ribosome components is required for proper G1/S transition in Saccharomyces cerevisiae. J Biol Chem 288: 31689–31700.

Gorenstein C, Warner JR. 1977. Synthesis and turnover of ribosomal proteins in the absence of 60S subunit assembly in *Saccharomyces cerevisiae*. Mol Gen Genet 157: 327–332.

Gregory B, Rahman N, Bommakanti A, Shamsuzzaman M, Thapa M, Lescure A, Zengel JM, Lindahl L. 2019. The small and large ribosomal subunits depend on each other for stability and accumulation. Life Sci Alliance 2.

Heinaniemi M, Nykter M, Kramer R, Wienecke-Baldacchino A, Sinkkonen L, Zhou JX, Kreisberg R, Kauffman SA, Huang S, Shmulevich I. 2013. Gene-pair expression signatures reveal lineage control. Nat Methods 10: 577–583.

Helser TL, Baan RA, Dahlberg AE. 1981. Characterization of a 40S ribosomal subunit complex in polyribosomes of Saccharomyces cerevisiae treated with cycloheximide. Mol Cell Biol 1: 51–57.

Hinnebusch AG, Ivanov IP, Sonenberg N. 2016. Translational control by 5'-untranslated regions of eukaryotic mRNAs. Science 352: 1413–1416.

Ho YH, Gasch AP. 2015. Exploiting the yeast stress-activated signaling network to inform on stress biology and disease signaling. Current genetics 61: 503–511.

Huber A, French SL, Tekotte H, Yerlikaya S, Stahl M, Perepelkina MP, Tyers M, Rougemont J, Beyer AL, Loewith R. 2011. Sch9 regulates ribosome biogenesis via Stb3, Dot6 and Tod6 and the histone deacetylase complex RPD3L. EMBO J 30: 3052–3064.

Jain A, Nilatawong P, Mamak N, Jensen LT, Jensen AN. 2020. Disruption in iron homeostasis and impaired activity of iron-sulfur cluster containing proteins in the yeast model of Shwachman-Diamond syndrome. Cell Biosci 10: 105.

James A, Wang Y, Raje H, Rosby R, DiMario P. 2014. Nucleolar stress with and without p53. Nucleus 5: 402–426.

Jung US, Sobering AK, Romeo MJ, Levin DE. 2002. Regulation of the yeast Rlm1 transcription factor by the Mpk1 cell wall integrity MAP kinase. Mol Microbiol 46: 781–789.

Juszkiewicz S, Slodkowicz G, Lin Z, Freire-Pritchett P, Peak-Chew SY, Hegde RS. 2020. Ribosome collisions trigger cis-acting feedback inhibition of translation initiation. Elife 9.

Kamburov A, Pentchev K, Galicka H, Wierling C, Lehrach H, Herwig R. 2011. ConsensusPathDB: toward a more complete picture of cell biology. Nucleic Acids Res 39: D712–717.

Kief DR, Warner JR. 1981. Coordinate control of syntheses of ribosomal ribonucleic acid and ribosomal proteins during nutritional shift-up in Saccharomyces cerevisiae. Mol Cell Biol 1: 1007–1015.

Kim D, Pertea G, Trapnell C, Pimentel H, Kelley R, Salzberg SL. 2013. TopHat2: accurate alignment of transcriptomes in the presence of insertions, deletions and gene fusions. Genome biology 14: R36.

Kim KY, Truman AW, Caesar S, Schlenstedt G, Levin DE. 2010. Yeast Mpk1 cell wall integrity mitogen-activated protein kinase regulates nucleocytoplasmic shuttling of the Swi6 transcriptional regulator. Mol Biol Cell 21: 1609–1619.

Kjeldgaard NO, Maaloe O, Schaechter M. 1958. The transition between different physiological states during balanced growth of Salmonella typhimurium. J Gen Microbiol 19: 607–616.

Klinge S, Woolford JL, Jr. 2019. Ribosome assembly coming into focus. Nat Rev Mol Cell Biol 20: 116–1

Kofler L, Prattes M, Bergler H. 2020. From Snapshots to Flipbook-Resolving the Dynamics of Ribosome Biogenesis with Chemical Probes. Int J Mol Sci 21.

Kondrashov N, Pusic A, Stumpf CR, Shimizu K, Hsieh AC, Xue S, Ishijima J, Shiroishi T, Barna M. 2011. Ribosome-mediated specificity in hox mRNA translation and vertebrate tissue patterning. Cell 145: 383–397.

Le Cao KA, Rohart F, McHugh L, Korn O, Wells CA. 2014. YuGene: a simple approach to scale gene expression data derived from different platforms for integrated analyses. Genomics 103: 239–251.

Li Y, Moir RD, Sethy-Coraci IK, Warner JR, Willis IM. 2000. Repression of ribosome and tRNA synthesis in secretion-defective cells is signaled by a novel branch of the cell integrity pathway. Mol Cell Biol 20: 3843–3851.

Liao Y, Smyth GK, Shi W. 2013. The Subread aligner: fast, accurate and scalable read mapping by seed-and-vote. Nucleic Acids Res 41: e108.

Love MI, Huber W, Anders S. 2014. Moderated estimation of fold change and dispersion for RNA-seq data with DESeq2. Genome biology 15: 550.

Lutfiyya LL, Johnston M. 1996. Two zinc-finger-containing repressors are responsible for glucose repression of SUC2 expression. Mol Cell Biol 16: 4790–4797.

MacDonald C, Payne JA, Aboian M, Smith W, Katzmann DJ, Piper RC. 2015. A family of tetraspans organizes cargo for sorting into multivesicular bodies. Dev Cell 33: 328–342.

Maitra N, He C, Blank HM, Tsuchiya M, Schilling B, Kaeberlein M, Aramayo R, Kennedy BK, Polymenis M. 2020. Translational control of one-carbon metabolism underpins ribosomal protein phenotypes in cell division and longevity. Elife 9.

Marion RM, Regev A, Segal E, Barash Y, Koller D, Friedman N, O’Shea EK. 2004. Sfp1 is a stress- and nutrient-sensitive regulator of ribosomal protein gene expression. Proc Natl Acad Sci U S A 101: 14315–14322.

Martin DE, Soulard A, Hall MN. 2004. TOR regulates ribosomal protein gene expression via PKA and the Forkhead transcription factor FHL1. Cell 119: 969–979.

Mills EW, Green R. 2017. Ribosomopathies: There’s strength in numbers. Science 358.

Mizuta K, Warner JR. 1994. Continued functioning of the secretory pathway is essential for ribosome synthesis. Mol Cell Biol 14: 2493–2502.

Monteiro PT, Mendes ND, Teixeira MC, d’Orey S, Tenreiro S, Mira NP, Pais H, Francisco AP, Carvalho AM, Lourenco AB et al. 2008. YEASTRACT-DISCOVERER: new tools to improve the analysis of transcriptional regulatory associations in Saccharomyces cerevisiae. Nucleic Acids Res 36: D132–136.

Nierras CR, Warner JR. 1999. Protein kinase C enables the regulatory circuit that connects membrane synthesis to ribosome synthesis in Saccharomyces cerevisiae. J Biol Chem 274: 13235–13241.

Niles BJ, Powers T. 2012. Plasma membrane proteins Slm1 and Slm2 mediate activation of the AGC kinase Ypk1 by TORC2 and sphingolipids in S. cerevisiae. Cell Cycle 11: 3745–3749.

Niles BJ, Powers T. 2014. TOR complex 2-Ypk1 signaling regulates actin polarization via reactive oxygen species. Mol Biol Cell 25: 3962–3972.

Oeffinger M, Tollervey D. 2003. Yeast Nop15p is an RNA-binding protein required for pre-rRNA processing and cytokinesis. Embo J 22: 6573–6583.

Ott M, Amunts A, Brown A. 2016. Organization and Regulation of Mitochondrial Protein Synthesis. Annu Rev Biochem 85: 77–101.

Pakos-Zebrucka K, Koryga I, Mnich K, Ljujic M, Samali A, Gorman AM. 2016. The integrated stress response. EMBO Rep 17: 1374–1395.

Piazzi M, Bavelloni A, Gallo A, Faenza I, Blalock WL. 2019. Signal Transduction in Ribosome Biogenesis: A Recipe to Avoid Disaster. Int J Mol Sci 20.

Pöll G, Braun T, Jakovljevic J, Neueder A, Jakob S, Woolford JL, Jr., Tschochner H, Milkereit P. 2009. rRNA maturation in yeast cells depleted of large ribosomal subunit proteins. PLoS One 4: e8249.

Polymenis M, Aramayo R. 2015. Translate to divide: сontrol of the cell cycle by protein synthesis. Microb Cell 2: 94–104.

Raiser DM, Narla A, Ebert BL. 2014. The emerging importance of ribosomal dysfunction in the pathogenesis of hematologic disorders. Leuk Lymphoma 55: 491–500.

Rousseau A, Bertolotti A. 2018. Regulation of proteasome assembly and activity in health and disease. Nat Rev Mol Cell Biol 19: 697–712.

Sanz M, Trilla JA, Duran A, Roncero C. 2002. Control of chitin synthesis through Shc1p, a functional homologue of Chs4p specifically induced during sporulation. Mol Microbiol 43: 1183–1195.

Shamsuzzaman M, Bommakanti A, Zapinsky A, Rahman N, Pascual C, Lindahl L. 2017. Analysis of cell cycle parameters during the transition from unhindered growth to ribosomal and translational stress conditions. PLoS One 12: e0186494.

Singh J, Tyers M. 2009. A Rab escort protein integrates the secretion system with TOR signaling and ribosome biogenesis. Genes Dev 23: 1944–1958.

Smith SJ, Crowley JH, Parks LW. 1996. Transcriptional regulation by ergosterol in the yeast Saccharomyces cerevisiae. Mol Cell Biol 16: 5427–5432.

Sung MK, Porras-Yakushi TR, Reitsma JM, Huber FM, Sweredoski MJ, Hoelz A, Hess S, Deshaies RJ. 2016. A conserved quality-control pathway that mediates degradation of unassembled ribosomal proteins. Elife 5.

Thapa M, Bommakanti A, Shamsuzzaman M, Gregory B, Samsel L, Zengel JM, Lindahl L. 2013. Repressed synthesis of ribosomal proteins generates protein-specific cell cycle and morphological phenotypes. Mol Biol Cell 24: 3620–3633.

Ulery TL, Jang SH, Jaehning JA. 1994. Glucose repression of yeast mitochondrial transcription: kinetics of derepression and role of nuclear genes. Mol Cell Biol 14: 1160–1170.

Ulirsch JC, Verboon JM, Kazerounian S, Guo MH, Yuan D, Ludwig LS, Handsaker RE, Abdulhay NJ, Fiorini C, Genovese G et al. 2018. The Genetic Landscape of Diamond-Blackfan Anemia. Am J Hum Genet 103: 930–947.

Vik A, Rine J. 2001. Upc2p and Ecm22p, dual regulators of sterol biosynthesis in Saccharomyces cerevisiae. Mol Cell Biol 21: 6395–6405.

Vind AC, Snieckute G, Blasius M, Tiedje C, Krogh N, Bekker-Jensen DB, Andersen KL, Nordgaard C, Tollenaere MAX, Lund AH et al. 2020. ZAKalpha Recognizes Stalled Ribosomes through Partially Redundant Sensor Domains. Mol Cell 78: 700–713 e707.

Warren AJ. 2018. Molecular basis of the human ribosomopathy Shwachman-Diamond syndrome. Advances in biological regulation 67: 109–127.

Weiss EL. 2012. Mitotic exit and separation of mother and daughter cells. Genetics 192: 1165–1202.

Wu CC, Peterson A, Zinshteyn B, Regot S, Green R. 2020. Ribosome Collisions Trigger General Stress Responses to Regulate Cell Fate. Cell 182: 404–416 e414.

Wu CC, Zinshteyn B, Wehner KA, Green R. 2019. High-Resolution Ribosome Profiling Defines Discrete Ribosome Elongation States and Translational Regulation during Cellular Stress. Mol Cell 73: 959–970 e955.

Zhang N, Wu J, Oliver SG. 2009. Gis1 is required for transcriptional reprogramming of carbon metabolism and the stress response during transition into stationary phase in yeast. Microbiology 155: 1690–1698.

Zhang YQ, Gamarra S, Garcia-Effron G, Park S, Perlin DS, Rao R. 2010. Requirement for ergosterol in V-ATPase function underlies antifungal activity of azole drugs. PLoS Pathog 6: e1000939.

Zhou X, Liao WJ, Liao JM, Liao P, Lu H. 2015. Ribosomal proteins: functions beyond the ribosome. J Mol Cell Biol 7: 92–104.

